# Migration Restores Hybrid Incompatibility Driven By Mitochondrial-Nuclear Sexual Conflict

**DOI:** 10.1101/2021.02.23.432505

**Authors:** Manisha Munasinghe, Benjamin C. Haller, Andrew G. Clark

## Abstract

In the mitochondrial genome, sexual asymmetry in transmission allows the accumulation of male-harming mutations since selection acts only on the effect of the mutation in females. Called the “Mother’s Curse”, this phenomenon induces a selective pressure for nuclear variants that compensate for this reduction in male fitness. Previous work has demonstrated the existence of these interactions and their potential to act as Dobzhansky–Muller incompatibilities, contributing to reproductive isolation between populations. However, it is not clear how readily they would give rise to and sustain hybrid incompatibilities. Here, we use computer simulations in SLiM 3 to investigate the consequences of sexually antagonistic mitochondrial-nuclear interactions in a subdivided population. We consider distinct migration schemes and vary the chromosomal location, and consequently the transmission pattern, of nuclear restorers. Disrupting these co-evolved interactions results in less-fit males, skewing the sex ratio toward females. Restoration of male fitness depends on both the chromosomal location of nuclear restorer loci and the migration scheme. Our results show that these interactions may act as Dobzhansky–Muller incompatibilities, but their strength is not enough to drive population isolation. Overall, this model shows the varied ways in which populations can respond to migration’s disruption of co-evolved mitochondrial-nuclear interactions.

## Introduction

A fundamental question in evolutionary genetics, and biology more broadly, is how new species form and remain distinct (1,2). Dobzhansky (3) and Mayr (4) argued that speciation hinges on the evolution of reproductive isolation, which limits gene flow between populations thereby advancing the process of speciation. The ‘biological species concept’ formalized this idea and explicitly defined species as groups of interbreeding natural populations that are substantially reproductively isolated from other groups (2,4). Reproductive isolation develops as isolating barriers accumulate, which prevent members of different populations from mating or forming zygotes (prezygotic), or which act after fertilization if hybrids are incompatible (postzygotic) (5,6). Understanding the genetic basis of hybrid incompatibility (which encompasses hybrid inviability, sterility, or reduced fitness compared to the parental populations) consequently allows us to better understand the mechanics of speciation (2).

Bateson (7), Dobzhansky (8), and Muller (9–11) first detailed how hybrid incompatibility could emerge between two allopatric populations. Populations acquire unique mutations while geographically separated, which may confer an adaptive advantage in their local environment, or may be neutral. When populations reunite and hybridize, evolutionarily untested interactions between these newly acquired mutations are exposed and may result in reduced hybrid fitness. Several empirical examples show that negative epistatic interactions, dubbed Dobzhansky– Muller incompatibilities, can generate hybrid incompatibility and thereby contribute to reproductive isolation (12–14). Consequently, there is continued interest in predicting which specific genetic interactions are likely to generate Dobzhansky–Muller incompatibilities to aid in their identification.

The unique function and transmission of mitochondrial DNA make nuclear-mitochondrial interactions promising candidates for Dobzhansky–Muller incompatibilities (15– 17). The mitochondrial genome encodes the ribosomal and transfer RNA components of the mitochondrial translation system, as well as 13 protein subunits that play an essential role in the electron transport chain and ATP synthase (18). Approximately 1,500 nuclear genes produce proteins that interact with the mitochondrial genome to ensure proper energy production (18–21). Coordination between these genomes is essential, as improper mitochondrial function is associated with various pathological phenotypes (22–24). Coadaptation between these two genomes is already thought to contribute to speciation, since nuclear genes evolve to compensate for the elevated deleterious load in the mitochondrial genome resulting from Muller’s Ratchet (25–27).

Different inheritance modes between the mitochondrial genome and the nuclear genome, however, naturally beget intergenomic conflict. Exclusively maternal transmission of mtDNA means only selection in females is effective. Consequently, sex-specific or sexually antagonistic mutations that are neutral or advantageous in females but deleterious in males can easily spread through populations (28–33). Coined the “Mother’s Curse” by Gemmell et al., these male-harming mitochondrial mutations have been studied most in plants, where they prevent pollen production in otherwise hermaphroditic species (28,34–37). The accumulation of male-harming mutations places selective pressure on the nuclear genome to evolve variants that restore male fitness, termed “restorers”, at least partially counteracting the cost of these Mother’s Curse variants. The rapid generation and fixation of such nuclear restorer mutations has been thoroughly documented in plants (38–40).

The ultimate dynamics of these interactions depend on the chromosomal location of a nuclear restorer. In an XY sex-determining system with an equal sex ratio, autosomes spend equal time in both sexes, the X spends ⅔ of its time in females, and the Y spends all its time in males. This difference influences several evolutionary processes. The mutation rate of a genetic element is generally higher the more time it spends in males (41–43), and the effective population sizes of the X and Y are reduced compared to autosomes (to ¾ and ¼, respectively), magnifying the effects of drift (44,45). Hemizygosity of sex chromosomes in males dramatically impacts the effect of selection on allele frequency dynamics. Finally, sexual antagonism can select for specific chromosomal locations to minimize the deleterious cost of specific variants in one sex by minimizing the time spent in that sex (46,47). A male-advantageous, female-harming mutation, for example, would experience some level of negative selection if present on an autosome or X chromosome but would escape this if Y-linked. Similarly, a mutation that is male-advantageous, female-neutral should evolve most rapidly on chromosomes that spend more time in males. Ultimately, the select effect of a mutation and the amount of time it spends in each sex (which is determined by its chromosomal location) will determine its evolutionary trajectory.

Exploration into the chromosomal placement of nuclear genes that interact with the mitochondrial genome reveals a complex landscape. Adaptive interactions in both sexes should result in movement of the associated nuclear gene onto the X, while those that work to mitigate sexual conflict, such as those involved with mitochondrial Mother’s Curse variants, will likely move off the X (48). Despite the gene-poor nature of the Y, strict paternal transmission of the Y should favor the accumulation of male-beneficial mutations, such as nuclear restorers of Mother’s Curse variants (49). Theoretical comparisons between autosomes, X, and Y suggest that nuclear restorers on the Y most rapidly spread and fix within a single population, too (50). The length of heterochromatin blocks on the Y may act as a sink for transcription factors or chromatin regulators, affecting the distribution of these elements in the rest of the genome (51,52). The demonstrated regulatory role of the Y may act to offset the costs of Mother’s Curse variants, since genes exhibiting sex-specific sensitivity to mtDNA are overrepresented among genes sensitive to Y-chromosomal variation (53,54).

Direct identification of Mother’s Curse nuclear restorers, which would provide insight into their chromosomal placement, has been limited, due in part by the experimental difficulty of identifying these interactions. Empirical studies that attempt this often use hybrid lines to disrupt co-evolved mito-nuclear interactions and evaluate differences in male and female hybrid fitness. This approach, while effective, is laborious and may capture only a few of these interactions, which are often sensitive to environmental conditions and so may evade detection (37,55–57). Nevertheless, these studies do highlight the potential of mito-nuclear interactions as Dobzhansky–Muller incompatibilities. The contribution of Mother’s Curse variants towards Haldane’s rule, the observation that the fitness of heterogametic hybrids is typically lower than homogametic hybrids, is often overlooked. The interaction between Mother’s Curse variants and nuclear restorers in XY systems could influence the early stages of speciation, but theoretical work on how these interactions evolve has mostly focused on their evolution within a single population (58–63).

Here, we construct a theoretical framework for exploring the disruption of co-evolved mito-nuclear interactions in multiple populations, and observe the resulting dynamics in simulations. We limit ourselves to two allopatric populations of equal size, each fixed for a unique set of mitochondrial Mother’s Curse variants and their corresponding nuclear restorers. A given simulation considers one of three chromosomal locations for these nuclear restorers: autosomal, X-linked, or Y-linked. At the beginning of each simulation, we select one of four migration schemes (detailed below). This design allows us to monitor hybrid fitness as migration creates gene flow between the populations and to assess whether mitochondrial-nuclear interactions can act to keep populations isolated – a key step in the speciation process.

## Material and Methods

### Model Design

Our model represents two allopatric populations that have independently evolved unique mito-nuclear interactions that are disrupted by gene flow between them. We consider two diploid, dioecious sexual populations that are initially geographically isolated. We define two classes of genomic elements: mitochondrial and nuclear. The mitochondrial genome is exclusively maternally inherited and homoplasmic, allowing us to treat it as haploid. Nuclear genomic elements may represent either an autosome, X, or Y, with their associated transmission patterns and ploidy. We model biallelic mitochondrial Mother’s Curse variants, where the wild-type variant is neutral in both sexes and the mutant is advantageous in females but deleterious in males. For each mitochondrial Mother’s Curse locus, there is a corresponding biallelic restorer locus in the nuclear genome that fully restores fitness of males carrying that mutant Mother’s Curse variant without impacting female fitness. **Table 1** shows the possible genotypes and fitnesses for one interaction. We calculated each individual’s final fitness multiplicatively across all interactions.

**Table 1.** Genotypes and Fitnesses of Males and Females for a single interaction (1 mtDNA Mother’s Curse Locus : 1 Nuclear Restorer Locus) depends on Nuclear Restorer Chromosomal Location. *M/A/X/Y* and *m/a/x/y* represent the wild type and mutant Mother’s Curse and restorer alleles respectively. *s*_*f*_ represents the advantage given to females by the Mother’s Curse variant, while *s*_*m*_ represents the cost of this variant in males. We assume incomplete dominance (*d*=0.5) for autosomal nuclear restorers.

| Nuclear Restorer Location | Females |  | Males |  |
| --- | --- | --- | --- | --- |
|  | Genotype | Fitness | Genotype | Fitness |
| Autosome | $M-AA$ | 1 | $M-AA$ | 1 |
| | $M-Aa$ | 1 | $M-Aa$ | 1 |
| | $M-aa$ | 1 | $M-aa$ | 1 |
| | $m-AA$ | $1+s_f$ | $m-AA$ | $1-s_m$ |
| | $m-Aa$ | $1+s_f$ | $m-Aa$ | $1-s_m$ |
| | $m-aa$ | $1+s_f$ | $m-aa$ | 1 |
| X | $M-XX$ | 1 | $M-XX$ | 1 |
| | $M-Xx$ | 1 | $M-xY$ | 1 |
| | $M-xx$ | 1 | $m-XY$ | $1-s_m$ |
| | $m-XX$ | $1+s_f$ | $m-xY$ | 1 |
| | $m-Xx$ | $1+s_f$ | | |
| | $m-xx$ | $1+s_f$ | | |
| Y | $M-XX$ | 1 | $M-XY$ | 1 |
| | $m-XX$ | $1+s_f$ | $M-Xy$ | 1 |
| | | | $m-XY$ | $1-s_m$ |
| | | | $m-Xy$ | 1 |

Each population starts with a fixed set of 20 mitochondrial Mother’s Curse variants and 20 corresponding fixed nuclear restorers.

These sets are disjoint, such that we track 40 loci in each of the mitochondrial and nuclear genomic elements, for a total of 80 loci across both populations. We chose to model 20 Mother’s Curse variants and nuclear restorers because there are no strong *a priori* expectations about their number, and because this allowed us to explore many interactions without excessively long simulation runtimes. For simplicity, we do not model new mutations at any point. We allow recombination for autosomal and X-linked nuclear restorers, and assume these restorers are unlinked (*r* = 0.5). Since recombination does not occur within the mitochondrial genome or the Y, we assume no recombination in those genomic elements (*r* = 0.0). We then allow migration to disrupt these co-evolved interactions between mitochondrial Mother’s Curse and nuclear restorer variants, using four different migration schemes: continuous symmetric migration, one generation of symmetric migration, continuous asymmetric migration, and continuous sex-symmetric migration (**Fig. S1.b**). We track the frequency of both native and foreign mutations over time to compare the degree of isolation between populations, defined in terms of the frequency of variants in both populations. We expect variants to be fixed in their population of origin and absent in the other under full isolation or found at comparable frequencies between the two populations if there is sufficient gene flow.

### SLiM Model Implementation

For these simulations, we use SLiM (version 3.3), a powerful, flexible genetic simulation framework capable of incorporating these design elements (64). We employ the nonWF model type in SLiM since it provides more direct control of model details, including the generation of offspring, migration, and epistatic fitness calculations. There are two key aspects of nonWF models in SLiM we must stress: how fitness is evaluated, and how populations are regulated. Fitness, in nonWF SLiM models, influences survival, and, consequently, fitness represents absolute fitness. **Table 1** details the fitness effect of one mito-nuclear interaction. The final fitness of an individual is calculated multiplicatively across all mutations possessed by an individual. This final fitness represents the likelihood that any given individual will survive to maturity. We enforce discrete, non-overlapping generations by setting the fitness of non-newborns to 0, ensuring they do not survive. Sex ratio and population size in SLiM nonWF models are emergent, determined by the individuals born in each generation minus those that died due to selection. As a result, the sex ratio may fluctuate if fitness differs between the sexes. The populations can grow up to a specified carrying capacity, but stochastic fluctuations in both population size and sex ratio are expected. Ultimately, our simulations are designed to robustly represent the demography of natural populations. Full details of the SLiM model design can be found in the supplement (Appendix S1). All scripts are on GitHub (https://github.com/mam737/mito_nuclear_SLiMulations).

## Results

Each model starts with two allopatric populations that have evolved a unique and fixed set of mitochondrial Mother’s Curse variants and nuclear restorers. We consider four distinct migration schemes (continuous symmetric migration, a single generation pulse of migration, continuous asymmetric migration, and continuous sex-specific migration) and three chromosomal locations for nuclear restorers (autosomal, X-linked, or Y-linked). For each scenario, we can track how migration between these populations impacts male fitness, the sex ratio, and allele frequency trajectories over time. We discuss models with continuous symmetric migration and continuous asymmetric migration below, with discussion on the remaining two relegated to the supplement (Appendix S2,3)

### Continuous Symmetric Migration

When migration occurs, male fitness is reduced in both populations. The importance of this reduction is governed by the magnitude of the difference between the benefit of Mother’s Curse variants in females and their cost in males (*s*_*f*_ - | *s*_*m*_|). Disrupting these co-evolved interactions leads to less fit males; as a result, more males than females die, skewing the sex ratio toward females. For autosomal and X-linked restorers, fitness recovery relies on the spread of both sets of nuclear restorers (NC1 and NC2). Recombination allows individuals to obtain restorers from both populations, protecting males from the deleterious effects of most or all Mother’s Curse variants. As autosomal restorers spread, the fitness and sex ratio slowly recover to initial levels (**Fig. 1.a**). X-linked restorers behave nearly identically to autosomal restorers (**Fig. S2)**.

**Figure 1.**
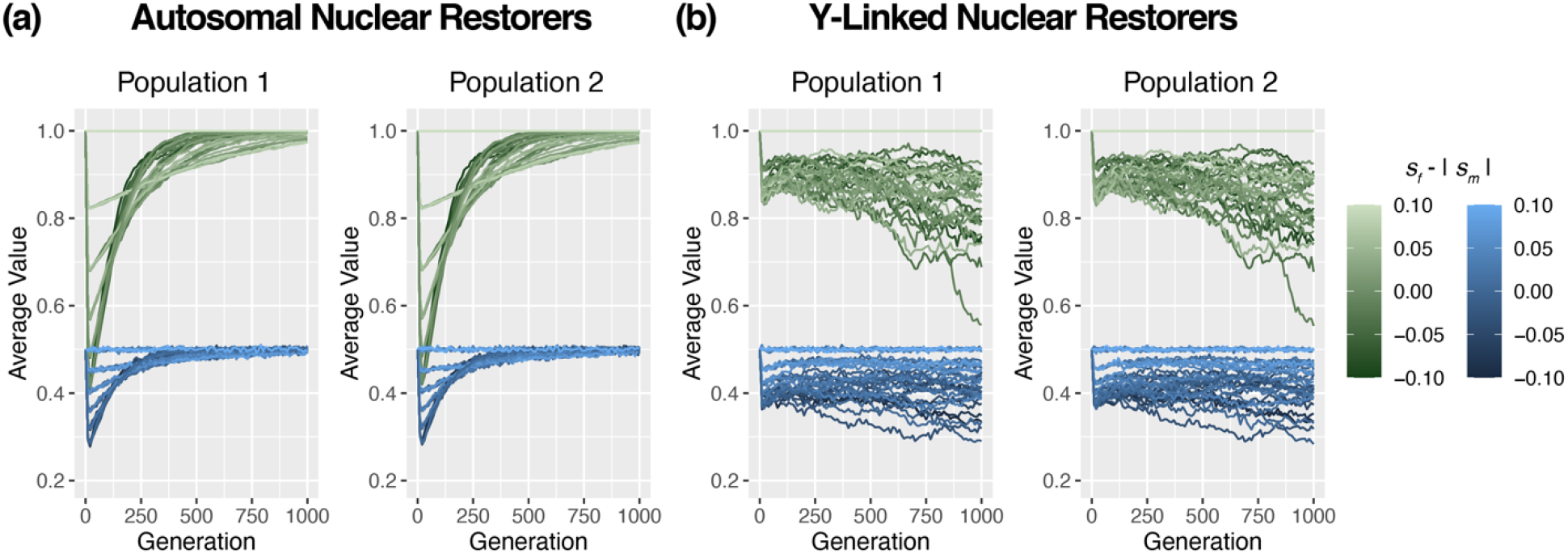
Mean male fitness (green) and sex ratio (blue) trajectories for population 1 (left) and population 2 (right) under continuous symmetric migration at rate *m* = 0.1. Simulation results for (a) autosomal nuclear restorers and (b) Y-linked restorers.

In contrast, Y-linked restorers are unable to recover male fitness. The absence of recombination on the Y makes it impossible for males to acquire both sets of nuclear restorers, and, consequently, hybrid males always suffer reduced fitness. However, it is worth noting that while Y-linked restorers suffer from a sustained reduction in male fitness, the size of this reduction is smaller in comparison with autosomal and X-linked restorers (**Fig. 1.b**). This is likely because all Y restorers from one population segregate together (another consequence of no recombination in the Y); as a result, a male that carries the Y haplotype matching their mitochondrial haplotype will be fully restored, while one that carries the other will experience the full cumulative cost of the Mother’s Curse variants.

We can assess the degree to which populations recover male fitness by examining the allele frequency trajectories of nuclear restorers for a specific *s*_*f*_ and *s*_*m*._ We find that the frequency of autosomal restorers tends toward fixation in both populations (**Fig. 2.a**, see **Fig. S3** for X-linked restorers). This aligns with both the restoration of male fitness and the return toward an equal sex ratio. The more deleterious the Mother’s Curse variants, the stronger the selective pressure is to obtain both sets of nuclear restorers. As more nuclear restorers fix, the deleterious effect of the mitochondrial haplogroup is minimized. The selective pressure for the remaining nuclear restorers is reduced, and their trajectories are determined primarily by drift. Once both sets of nuclear restorers are fixed, there is no fitness difference between the two mitochondrial haplotypes, and their frequency trajectory is determined by genetic drift thenceforth.

**Figure 2.**
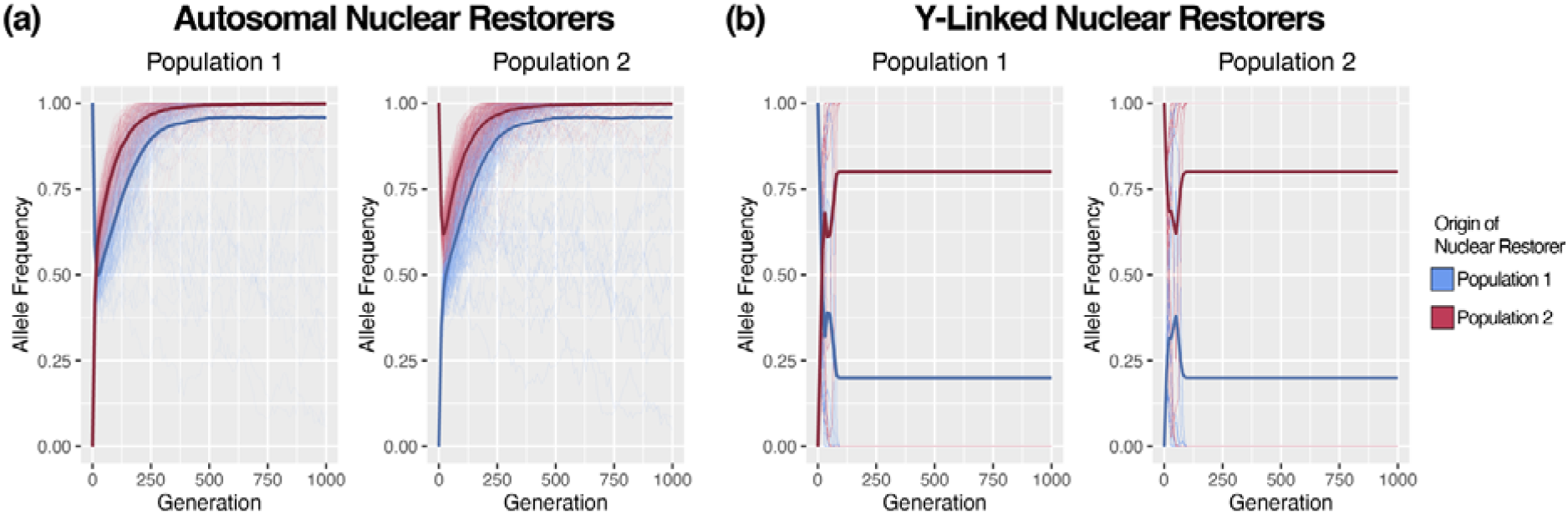
Allele Frequency Trajectories for both sets of nuclear restorers, those that originated in population 1 (blue) and population 2 (magenta) in each population (left and right) under continuous symmetric migration rate *m* = 0.1, *s*_*f*_ = 0.1, and *s*_*m*_ = -0.1. Each line represents the trajectory of a specific restorer. The darker, bold line represents the mean across all trajectories. (a) Shows the trajectories for our model with autosomal restorers, while (b) shows the trajectories for our model with Y-linked restorers.

With Y-linked restorers under the same conditions, we observe movement toward fixation of one Y haplotype and loss of the other; which haplotype is fixed versus lost seems to be initially stochastic, with positive feedback driving whichever haplotype initially increases in frequency toward fixation (**Fig. 2.b**).

There is no selection for either mitochondrial haplotype, since both haplotypes are equally fit in females. If one Y haplotype is lost from the population, male fitness for any individual carrying the associated mitochondrial haplogroup will always be reduced from that point on. Consequently, some males suffer reduced fitness depending on the frequency of the mitochondrial haplotypes. This explains why populations with autosomal or X-linked restorers show fitness recovery, while those with Y-linked restorers show significant variation in final fitness.

### Continuous Asymmetric Migration

For populations undergoing continuous asymmetric migration, we notice a distinct reduction in male fitness coupled with a skewed sex ratio as males die off, but the size and duration of this reduction depends on the migration rates between the two populations. For continuous asymmetric migration, we have two migration parameters, *m*_*1*_ and *m*_*2*_, where *m*_*1*_ determines the probability that an individual in population 1 will migrate into population 2, and *m*_*2*_ determines the reverse. We set *m*_*1*_ ≤ *m*_*2*_ such that population 1 receives more migrants from population 2 than the reverse. When the migration rates are asymmetric, we find that the sink population experiences a much more severe reduction in male fitness and a larger sex ratio skew. A return to the initial male fitness and sex ratio always occurs faster under continuous symmetric migration in comparison to continuous symmetric migration (∼150 generations to ∼500 generations respectively) with the time to recovery shrinking as the difference in migration rates increases. This is driven by variation in how male fitness is restored. Under continuous symmetric migration, we see the spread of both sets of nuclear restorers.

Under asymmetric migration, we instead see the domination of the MC2 and NR2 sets (initially associated with population 2). Both populations tend to rapidly fix for MC2 and NR2; however, occasionally the MC1 haplotype increases in frequency despite the asymmetric gene flow, which drives an increase in frequency of the NR1 set of restorers.

Continuous asymmetric migration with rates *m*_*1*_ = 0.01, *m*_*2*_ = 0.1 shows a smaller reduction in fitness and a shorter time to recovery than continuous symmetric migration with a rate of *m* = 0.1 (**Fig. 3**, see **Fig. S6** for X-linked restorers). This holds across all chromosomal locations for nuclear restorers. The allele frequency trajectories for both autosomal and Y-linked restorers show that population 1 becomes fixed for the mitochondrial haplotype (MC2) and nuclear restorers (NR2) present in population 2 (**Fig. 4**, see **Fig. S7** for X-linked restorers).

**Figure 3.**
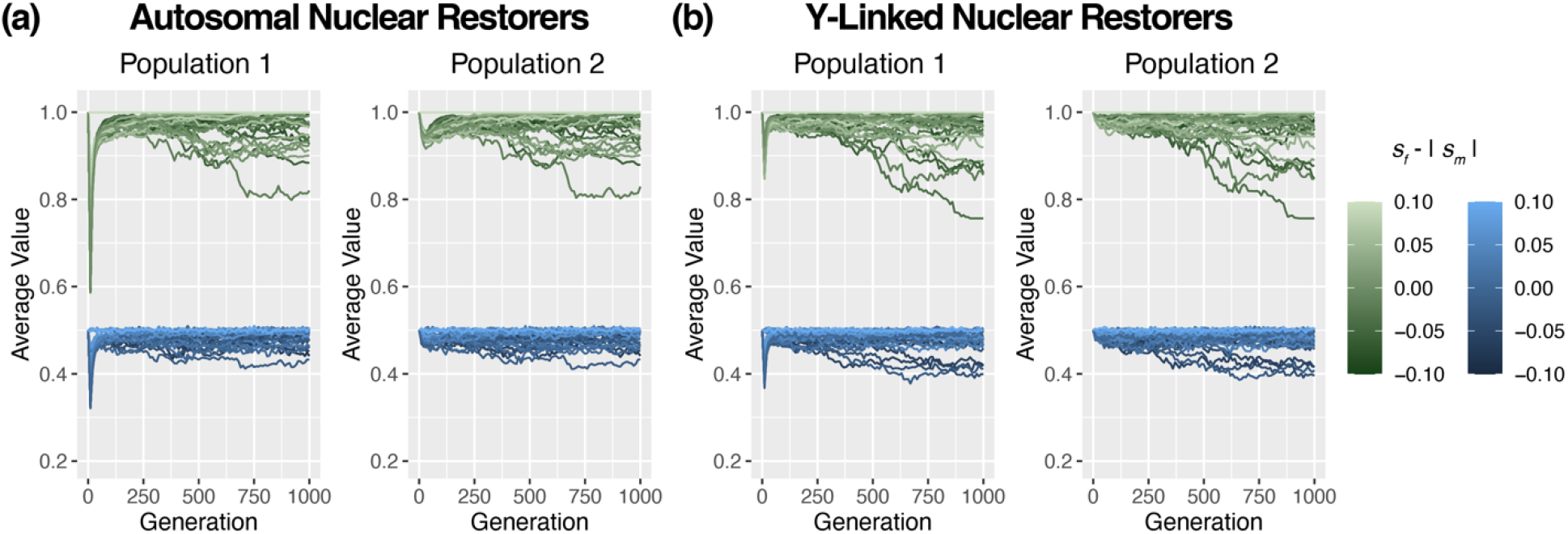
Mean male fitness (green) and sex ratio (blue) trajectories for population 1 (left) and population 2 (right) under continuous asymmetric migration rate *m*_*1*_ = 0.01 and *m*_*2*_ = 0.1. Simulation results for (a) autosomal nuclear restorers and (b) nuclear restorers.

**Figure 4.**
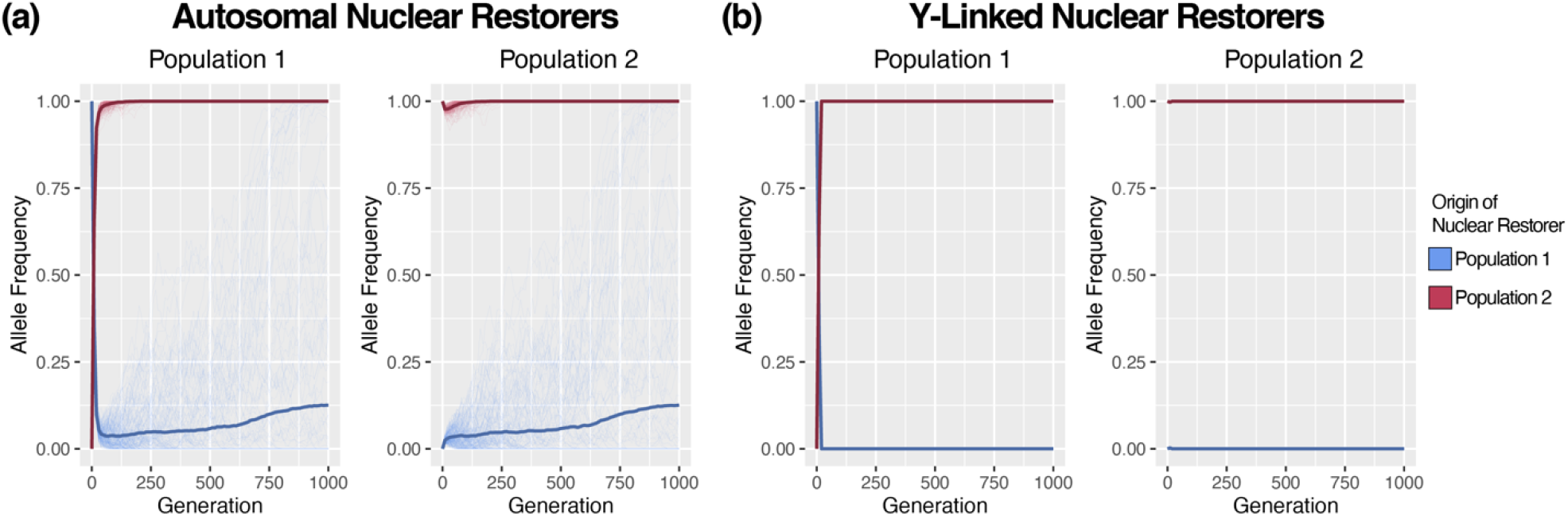
Allele Frequency Trajectories for both sets of nuclear restorers, those that originated in population 1 (blue) and population 2 (magenta) in each population (left and right) under continuous asymmetric migration rate *m*_*1*_ = 0.01, *m*_*2*_ = 0.1, *s*_*f*_ = 0.1, and *s*_*m*_ = -0.1. Each line represents the trajectory of a specific restorer. The darker, bold line represents the mean across all trajectories. (a) Shows the trajectories for our model with autosomal restorers, while (b) shows the trajectories for our model with Y-linked restorers.

Restoration relies on replacement due to the asymmetric gene flow, not on the spread of both sets of nuclear restorers, meaning there is little difference in the ability of autosomal, X-linked, or Y-linked restorers to rescue male fitness. The speed with which replacement occurs depends on *m*_*2*_, with larger *m*_*2*_ rates showing a smaller reduction in male fitness and faster time to restoration.

This occurs because a higher *m*_*2*_ expedites the replacement process; population replacement can occur more rapidly than recombination can merge the nuclear restorer sets.

As noted earlier, the NR1 set of nuclear restorers is not always lost. The fate of the NR1 set of restorers depends on whether the MC1 haplotype is able to increase in frequency. This is somewhat rare, since the MC1 haplotype’s frequency is also driven downward by the influx of migrants. Consequently, we more often see full population replacement, which may resemble older, simpler models of genetic drift under asymmetric gene flow.

## Discussion

Our results provide novel insights into the consequences of disrupting co-evolved mitochondrial-nuclear interactions. Continuous migration leads to a marked reduction in male fitness which skews the sex ratio as males die off. Populations respond to this in one of two ways, depending on whether the migration is symmetric or asymmetric. Under symmetric migration, populations acquire both sets of nuclear restorers to shield males from the deleterious effects of both mitochondrial haplotypes. Populations with Y-linked restorers are incapable of doing this, and they continue to suffer from reduced male fitness assuming both mitochondrial haplotypes remain common. However, the magnitude of this effect is mitigated, since all nuclear restorers for a specific Y haplotype segregate together; this means that any male with the corresponding mitochondrial haplotype is fully restored. Under asymmetric migration, one population’s genetic variation is usually replaced by the other due to gene flow, eliminating the potential for less-fit hybrid males. This occurs more rapidly than recombination and selection can merge the two sets of nuclear restorers, shortening the duration of reduced male fitness and the female-biased sex ratio in this scenario.

Under the parameters explored, we found little evidence that mito-nuclear interactions can lead to reproductive isolation. Disrupting these interactions generated less-fit male hybrids, implying that they do act as Dobzhansky–Muller incompatibilities, but this was not enough to keep populations isolated. Under continuous migration, populations responded to reduced male fitness either by incorporating all nuclear restorers through recombination, or by replacement after being swamped by gene flow. If we assumed that one mitochondrial haplogroup was more advantageous in females, the outcome would be analogous to population replacement since that haplotype would rapidly fix in both populations, consequently increasing the selective pressure for the associated set of nuclear restorers (Fig S12 and S13). However, even under extremely deleterious effect in males, such as male infertility which is one of the most commonly observed Mother’s Curse phenotypes, population isolation is not guaranteed as the spread of mitochondrial haplogroups is determined by their fitness effect in females (see Fig S14 and S15) (28,37,57,65).

Along with reduced male fitness, we see a distinct skew in the sex ratio as less-fit males are removed from the population. A 1:1 sex ratio is not a universal trait, even among dioecious species (66,67), but deviations from this have their own consequences: skewed sex ratios reduce the effective population size, which decreases the efficacy of selection (68). This increases the rates of genetic drift and inbreeding, ultimately resulting in decreased genetic variability (69). Over long periods of time, a significantly reduced number of males, as we observed in some of our simulations (up to 1:3 male:female), is likely to affect the genetic diversity of the Y even if nuclear restorers are autosomal or X-linked. Reduced male population sizes will affect the Y chromosome more strongly than autosomes or the X chromosome due to their exclusively male inheritance. The magnitude of the effect of reduced male population size on Y chromosome diversity is dependent on the initial population size. This may have large fitness consequences, since Y chromosomes influence a wide variety of traits (54,70).

Previous work has proposed the Y chromosome as a promising site for nuclear restorers of Mother’s Curse variants (50,71,72). Early empirical evidence by Innocenti et al. (73) and Rogell et al. (53) examined genes that are male-biased in their sensitivity to mitochondrial variation and found that these same genes are over-represented on a list of genes known to be sensitive to Y regulation, which suggests interactions between these two elements may influence male fitness. However, we show that under continuous symmetric migration, Y-linked restorers are outperformed by their autosomal or X-linked counterparts; the lack of recombination on the Y hinders a population’s ability to respond in the face of continual disruption of co-evolved mito-nuclear interactions. Parapatric populations with limited gene flow may regain fitness more rapidly with autosomal and X-linked restorers than with Y-linked restorers when contact with other populations leads to hybrid offspring. In a single population with no extrinsic gene flow, Y-linked nuclear restorers are the most effective since they spread throughout the population most rapidly (50). However, our results show that this does not hold if there is gene flow between two populations each experiencing the Mother’s Curse. Consequently, potential advantages of Y-linked restorers may only be realized in allopatric populations, for which there is little to no extrinsic gene flow.

Our theoretical framework and simulations explore the effects of migration and chromosomal location of nuclear restorers on mitochondrial-nuclear interactions. Among the many unexplored aspects of our model, several merit further research. We did not model the emergence of novel mitochondrial Mother’s Curse variants and nuclear restorers, nor the consequences of linkage for autosomal and X-linked restorers – mostly due to our inability to find robust empirical estimates of the relevant parameters. We also assumed that both populations were initially fixed for these interactions, consistent with theory (50,74–76), but the evolutionary dynamics of these mutations while they are still segregating may be of interest. We also did not explore the effects of genetic disequilibria between mitochondrial and nuclear genomic elements. It is well-established that several evolutionary forces, including genetic drift, epistatic selection, and nonrandom mating, may cause cytonuclear linkage disequilibria (i.e., departures from random association between nuclear and cytoplasmic genotypes) (74,75,77). Hybrid zones with directional and strong assortative mating will exacerbate cytonuclear disequilibria and epistatic interactions, like those we explored, and may further increase this non-random association between nuclear and cytoplasmic genotypes.

There are also facets of both Mother’s Curse variants and nuclear restorers that may influence our results. Mitochondrial DNA copy number ranges from hundreds to thousands of copies per cell depending on the cell’s energetic needs (78). If these copies are identical, they are “homoplasmic” and can be treated as a haploid element (which we assume). However, if there is variation among the mtDNA copies in a cell, which there often is, fitness is driven by more than the presence or absence of a specific variant; rather, the phenotype is determined by the proportion of mutant mtDNA (79). We also assume exclusive maternal inheritance of the mitochondrial genome, since only a handful of examples exist of paternal mitochondrial inheritance (whether partial or full) (80–82). Paternal transmission would introduce purifying selection in males on the male-deleterious Mother’s Curse mutations, but this is likely rare.

We also assume that a nuclear restorer is able to fully restore male fitness for its complementary mitochondrial Mother’s Curse variant, and that restorer mismatch (i.e., a negative fitness interaction between a nuclear restorer and the wild-type mitochondrial variant) does not occur. It is likely that restorer mismatch does exist in natural populations (see (83) for details on the differential strength of two nuclear restorers in *Brassica napus*), and it is our belief that this would influence the dynamics of our model. Our model makes several assumptions about the evolution of mitochondrial-nuclear interactions and possible migration patterns. It is possible different assumptions could lead to population isolation (e.g., longer bursts but not continuous migration, non-disjoint sets of Mother’s Curse variants and nuclear restorers, partial linkage of nuclear restorers, environmentally-dependent fitness evaluations, etc.). Additionally, even Dobzhansky–Muller incompatibilities that cannot produce complete isolation on their own may be important for isolation and speciation when acting in conjunction with other isolating factors (2).

Mother’s Curse variants demonstrably reduce the fitness of the heterogametic sex in XY sex-determining systems and the homogametic sex in ZW systems. Consequently, Mother’s Curse variants could possibly contribute to evidence of Haldane’s Rule in XY systems but not in ZW systems. Turelli and Moyle (2007) modeled the evolution of reciprocal hybrid asymmetry and clearly demonstrated a potential role for mitochondrial-nuclear interactions in contributing to asymmetric postmating reproductive isolation (85). However, we are unaware of any attempts to show an elevated presence of Haldane’s Rule or asymmetric reproductive isolation in XY-systems compared to ZW-systems. Such an endeavor could provide further insight into the influence of mito-nuclear interactions on reproductive isolation and speciation.

Finally, it is worth noting that selection on mtDNA variants can be influenced by their effect in males when variation in male and female fitness are directly correlated. Both inbreeding and kin selection couple a male’s fitness with that of his female relatives (62,63). The ability to fertilize a female is a male phenotype that directly affects a female’s reproductive success. If females mate with their male relatives, their reproductive success will be negatively impacted by the transmission of Mother’s Curse variants, even if those variants do not directly impact females. The evolution of the mitochondrial genome can consequently be influenced by the selective effect of variants in males if the connection between male and female reproductive success is sufficiently strong. Positive correlations between male and female fitness may select against male-harming variants, while negative correlations may select for these variants (84). The evolution of both Mother’s Curse variants and nuclear restorers will be influenced by these phenomena and may impact our results when inbreeding, kin selection, or mating that is dis(assortative) by mtDNA variants are present.

The model presented here provides novel insight into how populations respond to the disruption of co-evolved mitochondrial-nuclear interactions by migration, and it highlights the distinct forms this response takes depending on both the migration scheme and the chromosomal placement of nuclear restorers. Extensions of our model will provide additional insight into how asymmetrically inherited genomic elements can both cause and resolve genetic and sexual conflict.

## Supporting information

Full Supplemental Document

## Acknowledgements

We would like to thank Philipp Messer, Ian Vasconcellas Caldas, Mitchell Lokey, and Tram Nguyen for their assistance in testing and troubleshooting our SLiM models. Thanks also to Elissa Cosgrove for her assistance with several computational tasks.

## Author Contributions

M.M. and A.G.C. conceived the study and designed the theoretical framework. M.M. and B.H. incorporated the framework into SLiM and wrote all associated scripts. M.M. analyzed and visualized the results. M.M. wrote the first draft and all authors contributed to the writing of the manuscript. A.G.C. supervised the project.

## Data Accessibility

The scripts for all simulations can be found on GitHub:

https://github.com/mam737/mito_nuclear_SLiMulations

## References

1. Mayr E. Species, classification, and evolution. In: Biodiversity and Evolution. Tokyo: National Science Museum Foundation; 1995. p. 3–12.

2. Coyne J, Orr HA. Speciation. Sunderland, MA: Sinauer Associaties; 2004.

3. Dobzhansky T. A Critique of the Species Concept in Biology. Philos Sci. 1935;2(3):344–55.

4. Mayr E. Systematics and the Origin of Species. New York: Columbia University Press; 1942.

5. Dobzhansky T. Genetic Nature of Species Differences. Am Nat. 1937 Jul 1;71(735):404–20.

6. Dobzhansky T. Genetics and the Origins of Species. Third. New York: Columbia University Press; 1951.

7. Bateson W. Heredity and variation in modern lights. In: Darwin and Modern Science. Cambridge University Press; 1909. p. 85–101.

8. Dobzhansky T. Studies on hybrid sterility. Z Für Zellforsch Mikrosk Anat. 1934 Jan 1;21(2):169–223.

9. Muller HJ. Reversibility in Evolution Considered from the Standpoint of Genetics1. Biol Rev. 1939;14(3):261–80.

10. Muller HJ. Bearing of the Drosophila work on systematics. In: The New Systematics. Oxford: Clarendon Press; 1940. p. 185–268.

11. Muller H. Isolating mechanisms, evolution, and temperature. Biol Smposium. 1942;6:71–125.

12. Sweigart AL, Fishman L, Willis JH. A simple genetic incompatibility causes hybrid male sterility in mimulus. Genetics. 2006;172(4):2465–79.

13. Lowry DB, Modliszewski JL, Wright KM, Wu CA, Willis JH. Review. The strength and genetic basis of reproductive isolating barriers in flowering plants. Philos Trans R Soc Lond B Biol Sci. 2008 Sep 27;363(1506):3009–21.

14. Presgraves DC. Darwin and the origin of interspecific genetic incompatibilities. Am Nat. 2010 Dec;176 Suppl 1:S45–60.

15. Burton RS, Barreto FS. A disproportionate role for mtDNA in Dobzhansky-Muller incompatibilities? Mol Ecol. 2012 Oct;21(20):4942–57.

16. Jy C, Jy L. The Red Queen in mitochondria: cyto-nuclear co-evolution, hybrid breakdown and human disease. Front Genet. 2015 May 19;6:187–187.

17. Hénault M, Landry CR. When nuclear-encoded proteins and mitochondrial RNAs do not get along, species split apart. EMBO Rep. 2017 Jan 1;18(1):8–10.

18. Calvo SE, Mootha VK. The Mitochondrial Proteome and Human Disease. Annu Rev Genomics Hum Genet. 2010;11:25–44.

19. Neupert W, Herrmann JM. Translocation of Proteins into Mitochondria. Annu Rev Biochem. 2007 Jun 7;76(1):723–49.

20. Gershoni M, Templeton AR, Mishmar D. Mitochondrial bioenergetics as a major motive force of speciation. BioEssays. 2009;31(6):642–50.

21. Schmidt O, Pfanner N, Meisinger C. Mitochondrial protein import: from proteomics to functional mechanisms. Nat Rev Mol Cell Biol. 2010 Sep;11(9):655–67.

22. Duchen MR. Mitochondria in health and disease: perspectives on a new mitochondrial biology. Mol Aspects Med. 2004 Aug 1;25(4):365–451.

23. Pieczenik SR, Neustadt J. Mitochondrial dysfunction and molecular pathways of disease. Exp Mol Pathol. 2007 Aug 1;83(1):84–92.

24. Gorman GS, Chinnery PF, DiMauro S, Hirano M, Koga Y, McFarland R, et al. Mitochondrial diseases. Nat Rev Dis Primer. 2016 Oct 20;2(1):1–22.

25. Muller HJ. The relation of recombination to mutational advance. Mutat Res Mol Mech Mutagen. 1964;1(1):2–9.

26. Hill GE. Mitonuclear coevolution as the genesis of speciation and the mitochondrial DNA barcode gap. Ecol Evol. 2016;6(16):5831–42.

27. Hill GE. Mitonuclear Compensatory Coevolution. Trends Genet. 2020 Jun 1;36(6):403–14.

28. Lewis D. Male Sterility in Natural Populations of Hermaphrodite Plants the Equilibrium Between Females and Hermaphrodites to Be Expected with Different Types of Inheritance. New Phytol. 1941;40(1):56–63.

29. Charlesworth B, Charlesworth D. A Model for the Evolution of Dioecy and Gynodioecy. Am Nat. 1978;112(988):975–97.

30. Frank SA. The Evolutionary Dynamics of Cytoplasmic Male Sterility. Am Nat. 1989;133(3):345–76.

31. Frank SA, Hurst LD. Mitochondria and male disease. Nature. 1996 Sep;383(6597):224–224.

32. Gemmell NJ, Metcalf VJ, Allendorf FW. Mother’s curse: the effect of mtDNA on individual fitness and population viability. Trends Ecol Evol. 2004 May 1;19(5):238–44.

33. Vaught RC, Dowling DK. Maternal inheritance of mitochondria: implications for male fertility? Reprod Camb Engl. 2018;155(4):R159–68.

34. Kaul M. Male Sterility in Higher Plants. Springer Science & Business Media; 1988.

35. Budar F, Pelletier G. Male sterility in plants: occurrence, determinism, significance and use. Comptes Rendus Académie Sci Sér III Sci Vie. 2001 Jul 1;324:543–50.

36. Budar F, Touzet P, De Paepe R. The nucleo-mitochondrial conflict in cytoplasmic male sterilities revisited. Genetica. 2003 Jan;117(1):3–16.

37. Case AL, Finseth FR, Barr CM, Fishman L. Selfish evolution of cytonuclear hybrid incompatibility in Mimulus. Proc R Soc B Biol Sci. 2016 Sep 14;283(1838):20161493.

38. Chase CD. Cytoplasmic male sterility: a window to the world of plant mitochondrial–nuclear interactions. Trends Genet. 2007 Feb 1;23(2):81–90.

39. Arakawa T, Kagami H, Katsuyama T, Kitazaki K, Kubo T. A Lineage-Specific Paralog of Oma1 Evolved into a Gene Family from Which a Suppressor of Male Sterility-Inducing Mitochondria Emerged in Plants. Genome Biol Evol. 2020 Dec 6;12(12):2314–27.

40. Postel Z, Touzet P. Cytonuclear Genetic Incompatibilities in Plant Speciation. Plants [Internet]. 2020 Apr 10 [cited 2021 Jun 1];9(4). Available from: https://www.ncbi.nlm.nih.gov/pmc/articles/PMC7238192/

41. Malcom CM, Wyckoff GJ, Lahn BT. Genic mutation rates in mammals: local similarity, chromosomal heterogeneity, and X-versus-autosome disparity. Mol Biol Evol. 2003 Oct;20(10):1633–41.

42. Wilson Sayres MA, Makova KD. Genome analyses substantiate male mutation bias in many species. BioEssays News Rev Mol Cell Dev Biol. 2011 Dec;33(12):938–45.

43. Kirkpatrick M, Hall DW. Male-biased mutation, sex linkage, and the rate of adaptive evolution. Evol Int J Org Evol. 2004 Feb;58(2):437–40.

44. Vicoso B, Charlesworth B. Evolution on the X chromosome: unusual patterns and processes. Nat Rev Genet. 2006 Aug;7(8):645–53.

45. Mank JE. Small but mighty: the evolutionary dynamics of W and Y sex chromosomes. Chromosome Res Int J Mol Supramol Evol Asp Chromosome Biol. 2012 Jan;20(1):21–33.

46. Mank JE. Sex chromosomes and the evolution of sexual dimorphism: lessons from the genome. Am Nat. 2009 Feb;173(2):141–50.

47. Gibson JR, Chippindale AK, Rice WR. The X chromosome is a hot spot for sexually antagonistic fitness variation. Proc Biol Sci. 2002 Mar 7;269(1490):499–505.

48. Drown DM, Preuss KM, Wade MJ. Evidence of a Paucity of Genes That Interact with the Mitochondrion on the X in Mammals. Genome Biol Evol. 2012;4(8):875–80.

49. Bachtrog D. Y-chromosome evolution: emerging insights into processes of Y-chromosome degeneration. Nat Rev Genet. 2013 Feb;14(2):113–24.

50. Ågren JA, Munasinghe M, Clark AG. Sexual conflict through mother’s curse and father’s curse. Theor Popul Biol. 2019 Oct 1;129:9–17.

51. Henikoff S. Dosage-dependent modification of position-effect variegation in Drosophla. BioEssays. 1996;18(5):401–9.

52. Francisco FO, Lemos B. How Do Y-Chromosomes Modulate Genome-Wide Epigenetic States: Genome Folding, Chromatin Sinks, and Gene Expression. J Genomics. 2014 May 1;2:94–103.

53. Rogell B, Dean R, Lemos B, Dowling DK. Mito-nuclear interactions as drivers of gene movement on and off the X-chromosome. BMC Genomics. 2014 May 2;15(1):330.

54. Lemos B, Araripe LO, Hartl DL. Polymorphic Y Chromosomes Harbor Cryptic Variation with Manifold Functional Consequences. Science. 2008 Jan 4;319(5859):91–3.

55. Dowling DK, Friberg U, Lindell J. Evolutionary implications of non-neutral mitochondrial genetic variation. Trends Ecol Evol. 2008 Oct 1;23(10):546–54.

56. Arnqvist G, Dowling DK, Eady P, Gay L, Tregenza T, Tuda M, et al. Genetic architecture of metabolic rate: environment specific epistasis between mitochondrial and nuclear genes in an insect. Evol Int J Org Evol. 2010 Dec;64(12):3354–63.

57. Patel MR, Miriyala GK, Littleton AJ, Yang H, Trinh K, Young JM, et al. A mitochondrial DNA hypomorph of cytochrome oxidase specifically impairs male fertility in Drosophila melanogaster. VijayRaghavan K, editor. eLife. 2016 Aug 2;5:e16923.

58. Gregorius HR, Ross MD. Selection with Gene-Cytoplasm Interactions. I. Maintenance of Cytoplasm Polymorphisms. Genetics. 1984 May 1;107(1):165–78.

59. Clark AG. Natural selection with nuclear and cytoplasmic transmission. II. Tests with Drosophila from diverse populations. Genetics. 1985 Sep;111(1):97–112.

60. Babcock CS, Asmussen MA. Effects of Differential Selection in the Sexes on Cytonuclear Polymorphism and Disequilibria. Genetics. 1996 Oct 1;144(2):839–53.

61. Rand DM, Clark AG, Kann LM. Sexually antagonistic cytonuclear fitness interactions in Drosophila melanogaster. Genetics. 2001 Sep;159(1):173–87.

62. Unckless RL, Herren JK. Population genetics of sexually antagonistic mitochondrial mutants under inbreeding. J Theor Biol. 2009 Sep 7;260(1):132–6.

63. Wade MJ, Brandvain Y. Reversing mother’s curse: selection on male mitochondrial fitness effects. Evol Int J Org Evol. 2009 Apr;63(4):1084–9.

64. Haller BC, Messer PW. SLiM 3: Forward Genetic Simulations Beyond the Wright–Fisher Model. Mol Biol Evol. 2019 Mar 1;36(3):632–7.

65. Smith S, Turbill C, Suchentrunk F. Introducing mother’s curse: low male fertility associated with an imported mtDNA haplotype in a captive colony of brown hares. Mol Ecol. 2010 Jan;19(1):36–43.

66. Barrett SCH, Yakimowski SB, Field DL, Pickup M. Ecological genetics of sex ratios in plant populations. Philos Trans R Soc Lond B Biol Sci. 2010 Aug 27;365(1552):2549–57.

67. Hamilton WD. Extraordinary Sex Ratios. Science. 1967 Apr 28;156(3774):477.

68. Sewall Wright. Breeding Structure of Populations in Relation to Speciation. Am Nat. 1940 May 1;74(752):232–48.

69. Charlesworth B. Effective population size and patterns of molecular evolution and variation. Nat Rev Genet. 2009 Mar;10(3):195–205.

70. Lemos B, Branco AT, Hartl DL. Epigenetic effects of polymorphic Y chromosomes modulate chromatin components, immune response, and sexual conflict. Proc Natl Acad Sci U S A. 2010 Sep 7;107(36):15826–31.

71. Yee WKW, Rogell B, Lemos B, Dowling DK. Intergenomic interactions between mitochondrial and Y-linked genes shape male mating patterns and fertility in Drosophila melanogaster. Evolution. 2015;69(11):2876–90.

72. Dean R, Lemos B, Dowling DK. Context-dependent effects of Y chromosome and mitochondrial haplotype on male locomotive activity in Drosophila melanogaster. J Evol Biol. 2015 Oct;28(10):1861–71.

73. Innocenti P, Morrow EH, Dowling DK. Experimental Evidence Supports a Sex-Specific Selective Sieve in Mitochondrial Genome Evolution. Science. 2011 May 13;332(6031):845–8.

74. Clark AG. Natural Selection with Nuclear and Cytoplasmic Transmission. I. a Deterministic Model. Genetics. 1984 Aug;107(4):679–701.

75. Arnold J, Asmussen MA, Avise JC. An epistatic mating system model can produce permanent cytonuclear disequilibria in a hybrid zone. Proc Natl Acad Sci. 1988 Mar 1;85(6):1893–6.

76. Connallon T, Camus MF, Morrow EH, Dowling DK. Coadaptation of mitochondrial and nuclear genes, and the cost of mother’s curse. Proc R Soc B Biol Sci. 2018 Jan 31;285(1871):20172257.

77. Asmussen MA, Arnold J, Avise JC. Definition and properties of disequilibrium statistics for associations between nuclear and cytoplasmic genotypes. Genetics. 1987 Apr;115(4):755–68.

78. Cavelier L, Jazin E, Jalonen P, Gyllensten U. MtDNA substitution rate and segregation of heteroplasmy in coding and noncoding regions. Hum Genet. 2000 Jul;107(1):45–50.

79. Rossignol R, Faustin B, Rocher C, Malgat M, Mazat J-P, Letellier T. Mitochondrial threshold effects. Biochem J. 2003 Mar 15;370(Pt 3):751–62.

80. Havey MJ. Predominant Paternal Transmission of the Mitochondrial Genome in Cucumber. J Hered. 1997 May 1;88(3):232–5.

81. Schwartz M, Vissing J. Paternal Inheritance of Mitochondrial DNA. N Engl J Med. 2002 Aug 22;347(8):576–80.

82. Wolff JN, Nafisinia M, Sutovsky P, Ballard JWO. Paternal transmission of mitochondrial DNA as an integral part of mitochondrial inheritance in metapopulations of Drosophila simulans. Heredity. 2013 Jan;110(1):57–62.

83. Montgomery BR, Bailey MF, Brown GG, Delph LF. Evaluation of the cost of restoration of male fertility in Brassica napus. Botany. 2014 Sep 19;92(11):847–53.

84. Keaney TA, Wong HWS, Dowling DK, Jones TM, Holman L. Sibling rivalry versus mother’s curse: can kin competition facilitate a response to selection on male mitochondria? Proc Biol Sci. 2020 Jul 8;287(1930):20200575.

85. Turelli M, Moyle LC. Asymmetric Postmating Isolation: Darwin’s Corollary to Haldane’s Rule. Genetics. 2007 Jun 1;176(2):1059–88.

86. Munasinghe M, Haller BC, Clark AC. Migration Restores Hybrid Incompatibility Driven By Nuclear-Mitochondrial Sexual Conflict. bioRxiv doi:10.1101/2021.02.23.432505

