## Supplementary material for "Migration Restores Hybrid Incompatibility Driven By Mitochondrial-Nuclear Sexual Conflict": Full Supplemental Document

**Appendix S1: SLiM Model Implementation Details**

We initialize our model by defining two genomic elements: one mitochondrial and one nuclear (an autosome, X chromosome, or Y chromosome). We set the mutation rate to 0 throughout; we simulate only the mutations that we introduce at the start of each run. The recombination rate is set to 0.0 for the mitochondrial genome and thee Y chromosome, and we assume no linkage (a recombination rate of 0.5) for autosomes and the X chromosome. We establish four mutation classes: mitochondrial Mother’s Curse variants originating in population 1 (MC1), nuclear restorer variants originating in population 1 (NR1), mitochondrial Mother’s Curse variants originating in population 2 (MC2), and nuclear restorer variants originating in population 2 (NR2). We evaluate their fitness as detailed in **Table 1.** Note that every Mother’s Curse variant provides the same benefit to females and cost to males (i.e., *s_f_* and *s_m_* are constant for all Mother’s Curse variants, in a given run). We then construct two populations initially sized at 2000 individuals (this will serve as the carrying capacity), with an equal number of males and females. We then place our mutations, such that all individuals in population 1 have 20 unique pairs of MC1 and NR1 variants, while all individuals in population 2 have 20 unique pairs of MC2 and NR2 variants; each population is initially fixed (never heterozygous, although sometimes haploid or hemizygous) for its set of variants. Each MC and NR variant carries a specific tag such that each Mother’s Curse variant has one corresponding nuclear restorer, originally in the same population, that compensates for that MC variant’s effect on male fitness. The initial state represents a co-evolved set of MC and NR variants in each population, resulting from evolution in allopatry over some long (unmodeled) stretch of time. Once our populations are established, we start the simulation and allow migration, representing secondary contact between these previously allopatric populations.

Within each generation, the creation of offspring is the first step and is specified within a *reproduction()* callback. 1000 individuals are subsampled within each population, agnostic to their sex, to serve as parents. Male/female pairs are chosen randomly with replacement from this pool to generate a single offspring, until 2000 offspring have been generated. This allows us to have an equal number of males and females serving as parents, and we generate more offspring than the number of parents needed so that there will be enough individuals remaining after viability selection to serve as parents. This holds for all of our models, except for two of our supplemental models (when we have different female fitness for mitochondrial haplogroups (Fig S12 and S13) and more extreme values for the deleterious effects of Mother’s Curse variants in males (Fig S14 and S15) where we occasionally get incomplete replicates due to there being fewer than 1000 males).

SLiM itself has no understanding of mitochondrial DNA and is designed to model diploid genetics, so to achieve a functionally haploid genomic element we must take additional steps. The mitochondrial genome contains a marker mutation (i.e., neutral and used only as a placeholder) that allows us to ensure the maternal transmission of the mitochondrial genomic element by checking for the presence of this mutation in the maternally inherited mitochondrial genomic element and its absence in the paternally inherited mitochondrial genomic element. After this check is made, we clear the paternally inherited mitochondrial genomic element of any mutations as a safeguard, to ensure that there are no mitochondrial mutations are ever paternally. This results in a functionally haploid mitochondrial genomic element that is inherited maternally. We employ a similar method for any model that uses a Y-linked nuclear restorer, but in reverse, to ensure that the Y chromosome is exclusively transmitted to males, clearing both nuclear genomic elements in females. This allows us to have two genomic elements with different ploidies and transmission patterns in the same model, which is essential to modeling mitochondrial-nuclear interactions in SLiM.

After reproduction as described above, SLiM calculates the fitness of all individuals. We specifically set up fitness callbacks() to determine the fitness of males, as the fitness of a Mother’s Curse variant in males depends upon the number of nuclear restorers present. As mentioned earlier, we do not allow any new mutations to emerge. We also do not allow any recombination within either the mitochondrial genomic element or the Y nuclear genomic element. If the nuclear genomic element represents either an autosome or an X chromosome, we set the recombination rate to 0.5 so that each restorer is fully unlinked from the others. Because there is no recombination in the mitochondrial genome, an individual will always have either the 20 MC1s from population 1 or the 20 MC2s from population 2; we can consider these as two unchanging haplotypes that derive from their respective populations. Note that under this design, female fitness is equal and constant in both populations as it depends only on the mitochondrial haplotype present, and each haplotype has the same fitness of (1 + *s_f_*)^20^ since each haplotype contains 20 Mother’s Curse variants. Male fitness varies since it depends on the number of nuclear restorers present. Mean population fitness, prior to any density-dependent effects on fitness, is determined by the mean fitness of males in each population and the sex ratio.

After calculating the fitness of all individuals SLiM applies viability selection, with each individual’s probability of survival equal to its fitness (clipped to a maximun of 1.0, representing guaranteed survival; females are always guaranteed to survive, given the fitness model described above with the exception of our supplemental model with different female fitness for each mitochondrial haplogroup). From the remaining individuals, we select individuals to migrate to the other population depending on the migration scheme. For continuous symmetric migration, we have one migration parameter *m* which defines the probability that an individual will migrate from one population to the other. We use the same migration parameter to implement a single-generation pulse of symmetric migration, but after that first generation, we set *m* to 0 to prevent further migration. For continuous asymmetric migration, we have two migration parameters, *m_1_* and *m_2_*, where *m_1_* determines the probability that an individual in population 1 will migrate into population 2, and *m_2_* determines the reverse. We set *m_1_* ≤ *m_2_* such that population 1 receives more migrants from population 2 than the reverse. Finally, for continuous sex-specific migration, we assign two migration parameters *m_f_* and *m_m_*. *m_f_* is the probability that a female individual will move from one population to the other, while *m_m_* is the probability that a male individual will do so. Note that when *m_f_* = *m_m_* we replicate continuous symmetric migration, so we are particularly concerned with when *m_f_* ≠ *m_m_* (see **Fig. S1.b** for visual representation of all migration schemes).

After migration, the generation cycle then starts again; this repeats for 1000 generations. We track the mean fitness trajectories of each population, the allele frequencies of all variants, and the sex ratio every 10 generations to discern how the populations respond over time to disruption of their co-evolved mitochondrial-nuclear interactions due to migrational gene flow from secondary contact, for a specific set of parameters (migration parameters, *s_f_*, and *s_m_*).

All migration parameters (*m* under continuous symmetric migration and single-generation symmetric migration, *m_1_* and *m_2_* such that *m_1_* ≤ *m_2_* under continuous asymmetric migration, and *m_f_* and *m_m_* under continuous sex-specific migration) range from 0.01 to 0.1 in increments of 0.018, providing six distinct values for each. *s_f_* ranges from 0.0 to 0.1 in increments of 0.02, and *s_m_* similarly ranges from −0.1 to 0.0 in increments of 0.02, similarly providing six distinct values for each. For each specific parameter set, we replicate the simulation 10 times. Therefore, a total of 6480 (3*6*6*6*10), 6480, 232680 (3*6*6*21*10), and 38880 (3*6*6*36*10) replicates were run in total for continuous symmetric migration, a single-generation pulse of symmetric migration, continuous asymmetric migration, and continuous sex-specific migration respectively, making 284520 runs all together.

Additionally, to test certain assumptions about our model, we incorporated 3 additional supplemental models under the framework used for continuous symmetric migration. The first involved varying the dominance of autosomal restorers *h* from 0.0 (fully dominant) to 1.0 (fully recessive) in increments of 0.2 for a total of 6 values for a subset of migration parameter values (0.01,0.05,0.1) for a total of 6480 (1*3*6*6*6*10) runs. We also tested what would happen if the mitochondrial haplogroup native to each population had different fitness effects in females (i.e., a *s_f_p1_* and *s_f_p2_* for Mother’s Curse variants that evolved in population 1 and 2 respectively such that *s_f_p1_* ≤ *s_f_p2_*). For this model, we must incorporate a density-dependent mortality to account for the differentially advantageous mitochondrial haplogroups. We only consider autosomal and Y-linked nuclear restorers, a subset of migration parameter values (0.01,0.1), and a subset of *s_m_* values (−0.1, −0.01,0.0), which results in a total of 2520 (2*2*3*21*10) runs. Finally, we consider an extension where we allow more extreme values of *s_m_* (particularly −0.5 and −1.0)*_._* We once again consider only autosomal and Y-linked nuclear restorers and a subset of migration parameter values (0.01,0.05, and 0.1), for a total of 720 (2*3*2*6*10) runs.

**Appendix S2: Results for A Single Generation Pulse of Symmetric Migration**

If symmetric migration is allowed for only one generation, we observe a much smaller reduction in fitness which quickly returns to the initial level. One generation of migration does create less-fit hybrid males, but this reduction is not sustained. These F1 hybrid males have greatly reduced fitness and a large number of them die before producing any offspring. Later-generation hybrids are predominantly the result of F1 hybrid females mating with males native to the population they are in, and the male offspring from these matings will also have greatly reduced fitness. Without new migrants to continue generating hybrids, the number of hybrids in both populations declines until there are few to no hybrids left. Consequently, we see only minor fluctuations in male mean fitness and sex ratio (**Fig. S4,5)**. This is consistent across all nuclear restorer locations. Under our parameter values, one generation of symmetric migration is simply not enough to allow less-fit hybrid males to persist. We did not simulate longer bursts of migration, but we expect that if enough hybrid males are generated the scenario would be similar to continuous migration: recombination would spread nuclear restorers to offset reduced male fitness.

The code for every model can be found on GitHub: https://github.com/mam737/mito_nuclear_SLiMulations.

**Appendix S3: Results for Continuous Sex-Specific Migration**

Here, migration rates are symmetric between the populations, but different for males versus females. This model behaves similarly to continuous symmetric migration with a rate equal to the average of the male and female migration rates. This suggests that symmetry in the migration rates may be more influential that the difference in migration rates between the sexes (i.e., the male and female migration rates may be different from each other but are the same between the two populations) (**Fig S8,9**). As with continuous symmetric migration, we see reduced male fitness and an associated skew in the sex ratio for autosomal and X-linked nuclear restorers; male fitness and the sex slowly recover as both sets of restorers spread to both populations. The larger the female migration rate is relative to the male migration rate, the more rapid the decline in fitness is. The final state of the populations is essentially the same: recovery, in the case of autosomal and X-linked restorers, and a sustained reduction in fitness, for Y-linked restorers.

**Appendix S4: Results for Varying Dominance of Autosomal Nuclear Restorers Under Continuous Symmetric Migration**

The nature of X-linked and Y-linked nuclear restorers is such that males can only carry one copy, and we assume that a single copy is sufficient to fully restore male fitness. However, the impact of dominance on the strength of nuclear restorers may influence the evolution of autosomal restorers, distinguishing them from the other two classes. Consequently, for our model of continuous symmetric migration, we tested whether varying the dominance would greatly impact our results. We added the parameter *h*, which ranged from 0 to 1 with *h* = 0 representing a fully dominant restorer and *h* = 1 representing a fully recessive restorer. It is worth noting that this is the opposite of how dominance is usually thought of, but it is simply a consequence of how we defined our fitness calculations. We found that the general trajectory of male fitness remained unchanged as we varied dominance (**Fig S10**), but the magnitude of the reduction to both male fitness and the corresponding skew to the sex ratio increased as we increased the recessivity of the nuclear restorer. The allele frequency trajectories largely seem unaffected by dominance, which suggests both sets of nuclear restorers spread and increase in frequency in each population (**Fig S11**).

**Appendix S5: Results for Assigning Different Fitness Effects in Females for Each Mitochondrial Haplogroup Under Continuous Symmetric Migration**

Our models assume that the two mitochondrial haplogroups are equally fit in females. Biologically speaking, it is much more likely that Mother’s Curse mutations between populations would have different effects in females, and these effects may even be environment-specific. However, if we ignore environmentally-dependent fitness effects, migration between two populations where one mitochondrial haplotype is more fit in females than the other should result in an outcome analogous to population replacement. To confirm this, we did test what would happen in our models if we incorporated this. As expected, we observed a very rapid fixation of the mitochondrial haplotype with a higher female fitness. Females are consequently monomorphic, and we observe a rapid fixation of nuclear restorers associated with this haplotype in both populations **(Fig S13).** It is worth noting that this is the only model that incorporates density-dependence mortality in order to realize the differentially advantageous mitochondrial haplogroups, which explains why the fitness trajectories look dissimilar to other models **(Fig S12).**

**Appendix S6: Results for Extremely Deleterious Effects in Males Under Continuous Symmetric Migration**

We were curious whether more extreme values for *s_m_* would be sufficient to isolate populations. Even if we assume hybrid males are fully sterile, female transmission of the mitochondrial genome would still allow the mitochondrial haplogroup to invade. Continued migration would allow the offspring of hybrid females to obtain the necessary nuclear restorers (in the autosomal or X-linked case) such that their male offspring would experience full restoration. We tested this hypothesis by testing what would occur if we used *s_m_* values equal to −0.5 and −1.0 for Mother’s Curse variants. We found little evidence for population isolation, and, instead, we observed a very sharp reduction in male fitness and skew to the sex ratio that recovers very quickly (**Fig S14**). This is likely because males with mismatched mitochondrial haplogroups and nuclear restorers die during viability selection due to their extremely low fitness values. The allele frequency trajectories are very similar between both populations suggesting strong gene flow and sharing of restorers (**Fig S15)**. These results suggest that even extremely male-harming Mother’s Curse variants are not capable of resulting in population isolation on their own.

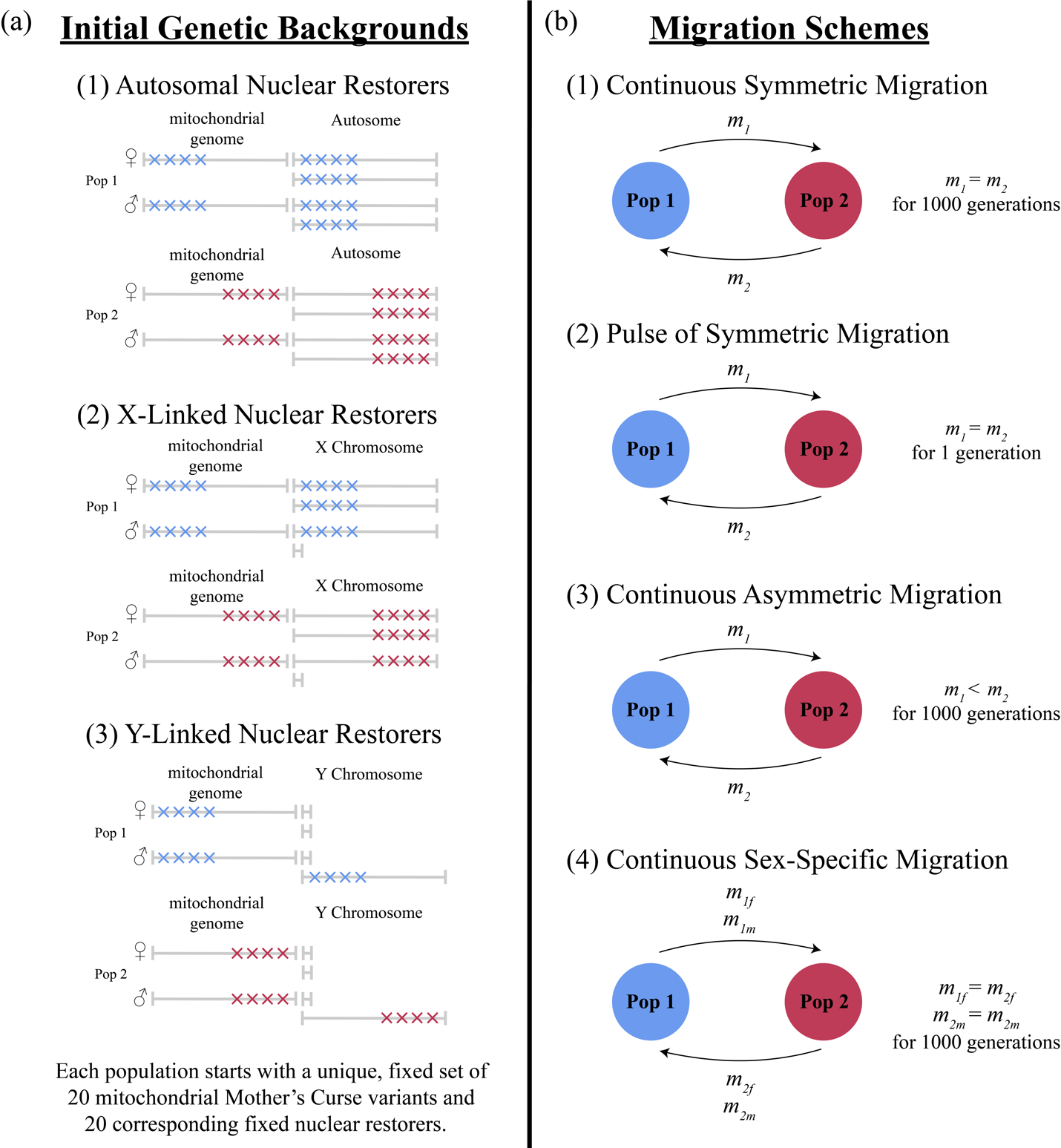

**Figure S1.** **Schematic of Model Design**. Blue represents population 1, while red represents population 2. (a) Representation of initial genetic backgrounds for females and males in each population, depicting the chromosomal location of Mother’s Curse variants and their associated nuclear restorers. Here, four of each is shown in each individual, whereas our simulations actually give 20 to each individual (b) Representation of the four distinct migration schemes explored.

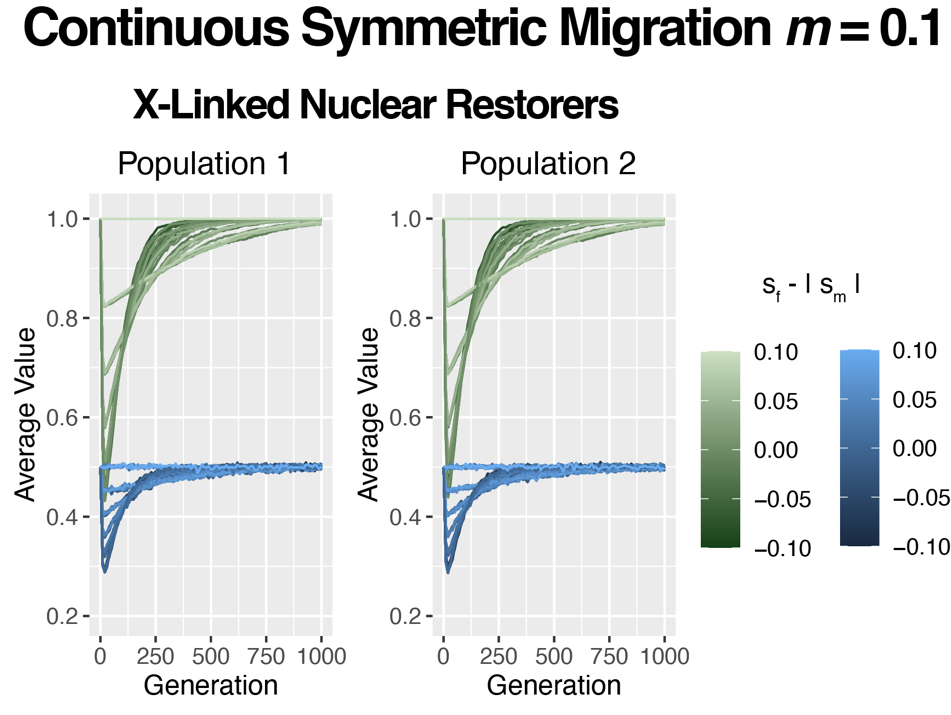

**Figure S2. Mean Male Fitness and Sex Ratio Trajectories for population 1 (left) and population 2 (right) under Continuous Symmetric Migration rate *m* = 0.1 for X-linked Nuclear Restorers.** Mean male fitness trajectories are colored in green, while the sex ratio is colored in blue. The specific color refers to the magnitude of difference between the advantage of Mother’s Curse variants in females and the deleterious effect of these variants in males.

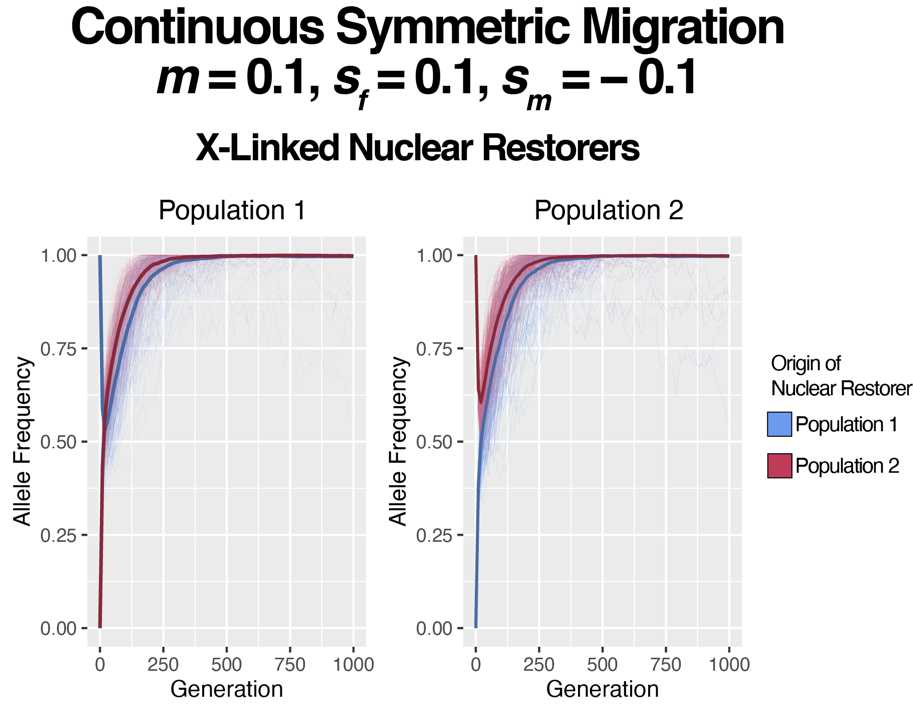

**Figure S3. Allele Frequency Trajectories for both sets of nuclear restorers, those that originated in population 1 (blue) and population 2 (magenta), in each population (left and right) under continuous symmetric migration rate *m* = 0.1, *s_f_* = 0.1, and *s_m_* = −0.1 for X-linked nuclear restorers**. Each line represents the trajectory of a specific restorer. The darker, bold line represents the mean across all trajectories for that class of nuclear restorers.

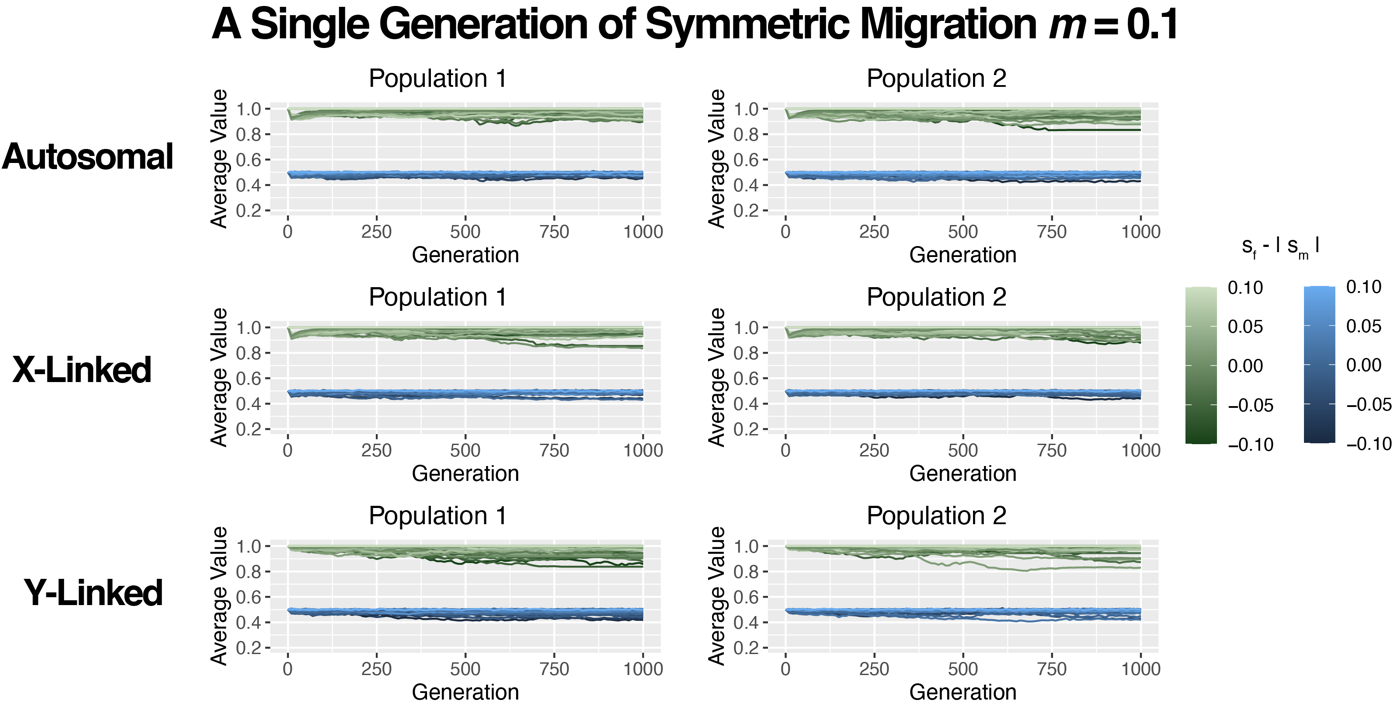

**Figure S4. Mean Male Fitness and Sex Ratio Trajectories for population 1 (left) and population 2 (right) under 1 Generation of Symmetric Migration rate *m* = 0.1 for Autosomal (Row 1), X-Linked (Row 2), and Y-Linked (Row 3) Restorers.** Mean male fitness trajectories are colored in green, while the sex ratio is colored in blue. The specific color refers to the magnitude of difference between the advantage of Mother’s Curse variants in females to the deleterious effect of these variants in males.

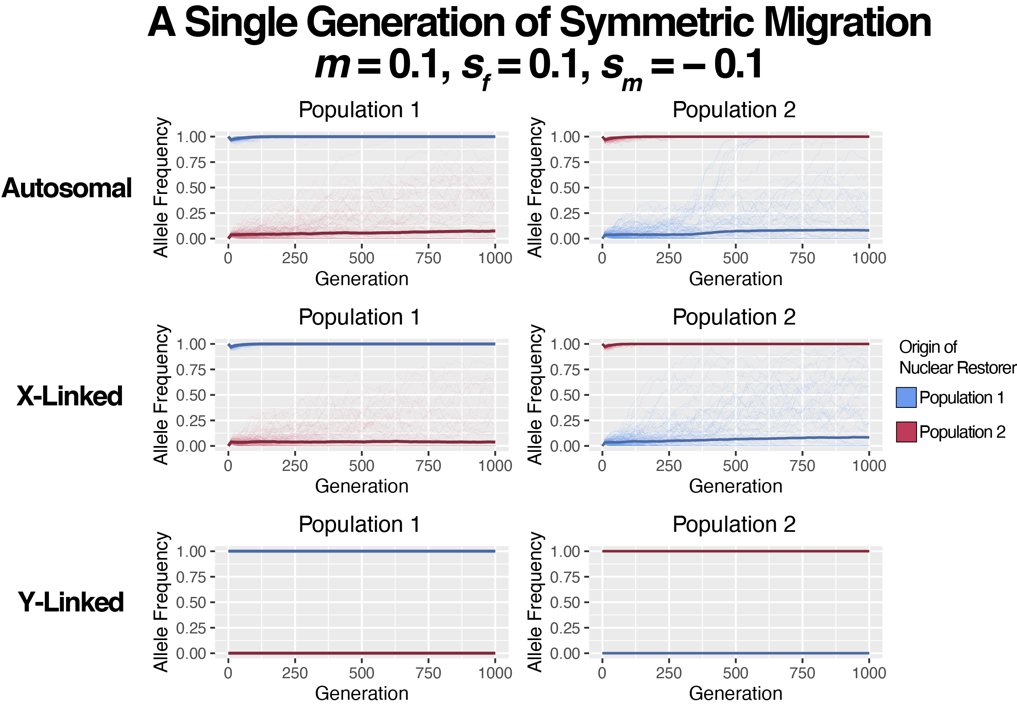

**Figure S5. Allele Frequency Trajectories for both sets of nuclear restorers, those that originated in population 1 (blue) and population 2 (magenta) in each population (left and right) under 1 generation of symmetric migration rate *m* = 0.1, *s_f_* = 0.1, and *s_m_* = −0.1**. Each line represents the trajectory of a specific restorer. The darker, bold line represents the mean across all trajectories for that class of nuclear restorers. The first row shows autosomal restorers, the second row shows X-linked restorers, and the third row shows Y-linked restorers.

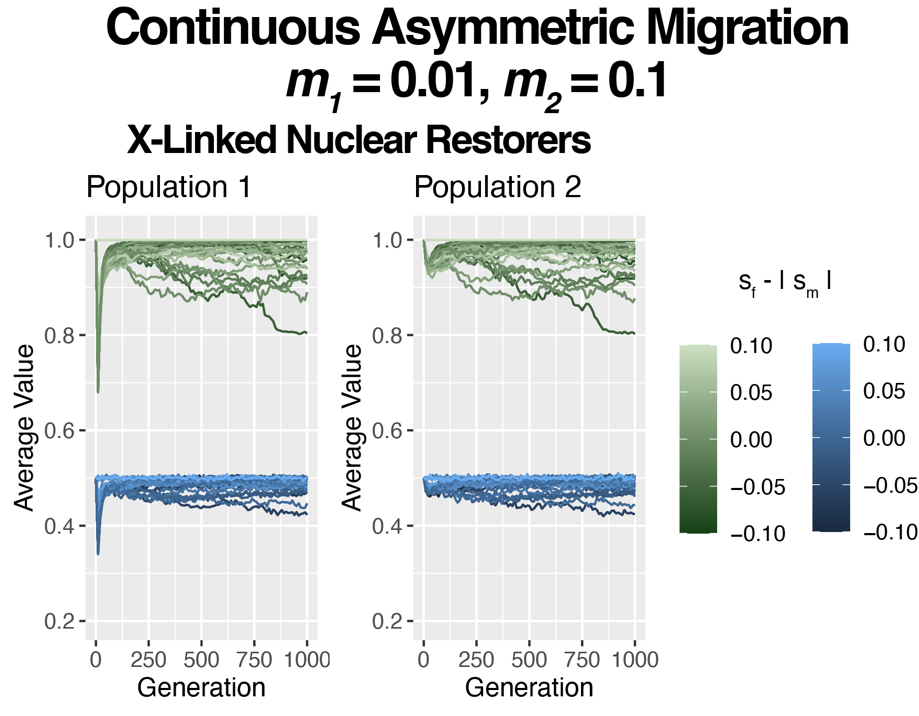

**Figure S6. Mean Male Fitness and Sex Ratio Trajectories for population 1 (left) and population 2 (right) under Continuous Asymmetric Migration rate *m_1_* = 0.01, *m_2_* = 0.1 for X-linked Nuclear Restorers.** Mean male fitness trajectories are colored in green, while the sex ratio is colored in blue. The specific color refers to the magnitude of difference between the advantage of Mother’s Curse variants in females and the deleterious effect of these variants in males.

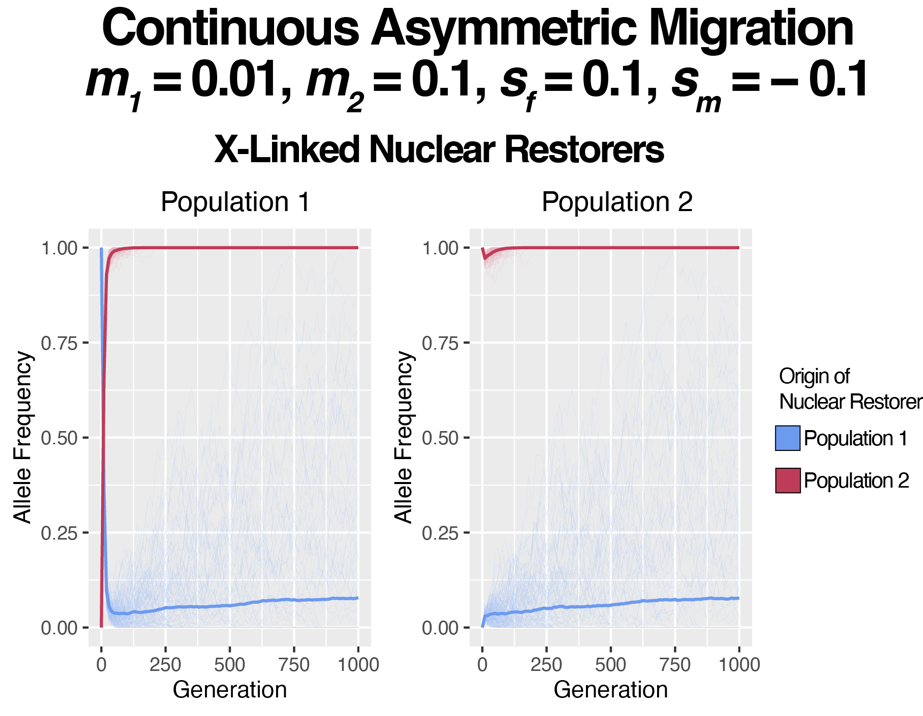

**Figure S7. Allele Frequency Trajectories for both sets of nuclear restorers, those that originated in population 1 (blue) and population 2 (magenta) in each population (left and right) under continuous asymmetric migration rate *m_1_* = 0.1, *m_2_* = 0.1, *s_f_* = 0.1, and *s_m_* = −0.1 for X-linked nuclear restorers**. Each line represents the trajectory of a specific restorer. The darker, bold line represents the mean across all trajectories for that class of nuclear restorers.

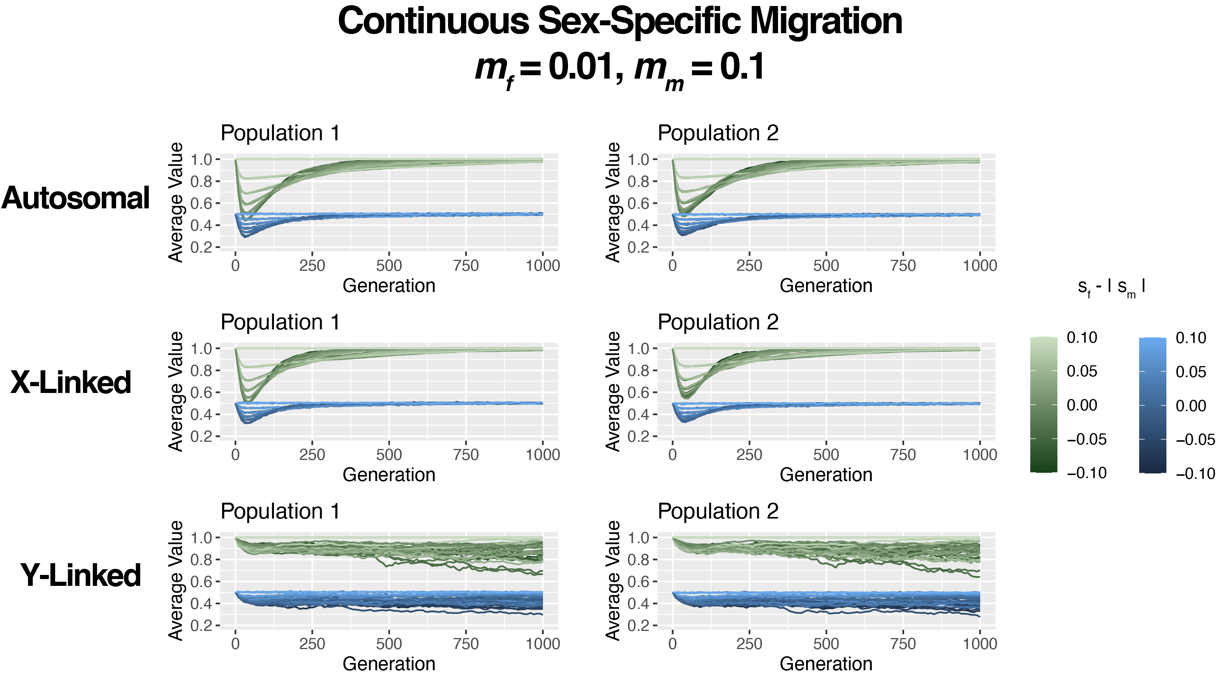

**Figure S8. Mean Male Fitness and Sex Ratio Trajectories for population 1 (left) and population 2 (right) under continuous sex-specific migration rates *m_f_* = 0.01, *m_m_* = 0.1 for Autosomal (Row 1), X-Linked (Row 2), and Y-Linked (Row 3) Restorers.** Mean male fitness trajectories are colored in green, while the sex ratio is colored in blue. The specific color refers to the magnitude of difference between the advantage of Mother’s Curse variants in females and the deleterious effect of these variants in males.

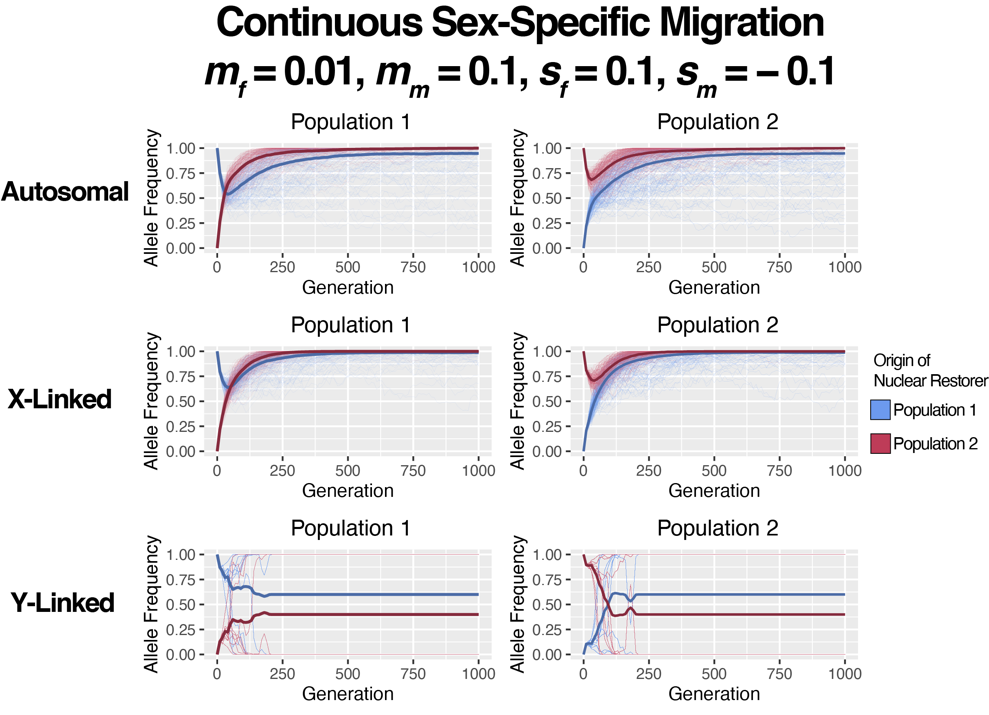

**Figure S9. Allele Frequency Trajectories for both sets of nuclear restorers, those that originated in population 1 (blue) and population 2 (magenta) in each population (left and right) continuous sex-specific migration rate *m_f_* = 0.01, *m_m_* = 0.1, *s_f_* = 0.1, and *s_m_* = −0.1**. Each line represents the trajectory of a specific restorer. The darker, bold line represents the mean across all trajectories for that class of nuclear restorers. The first row shows autosomal restorers, the second row shows X-linked restorers, and the third row shows Y-linked restorers.

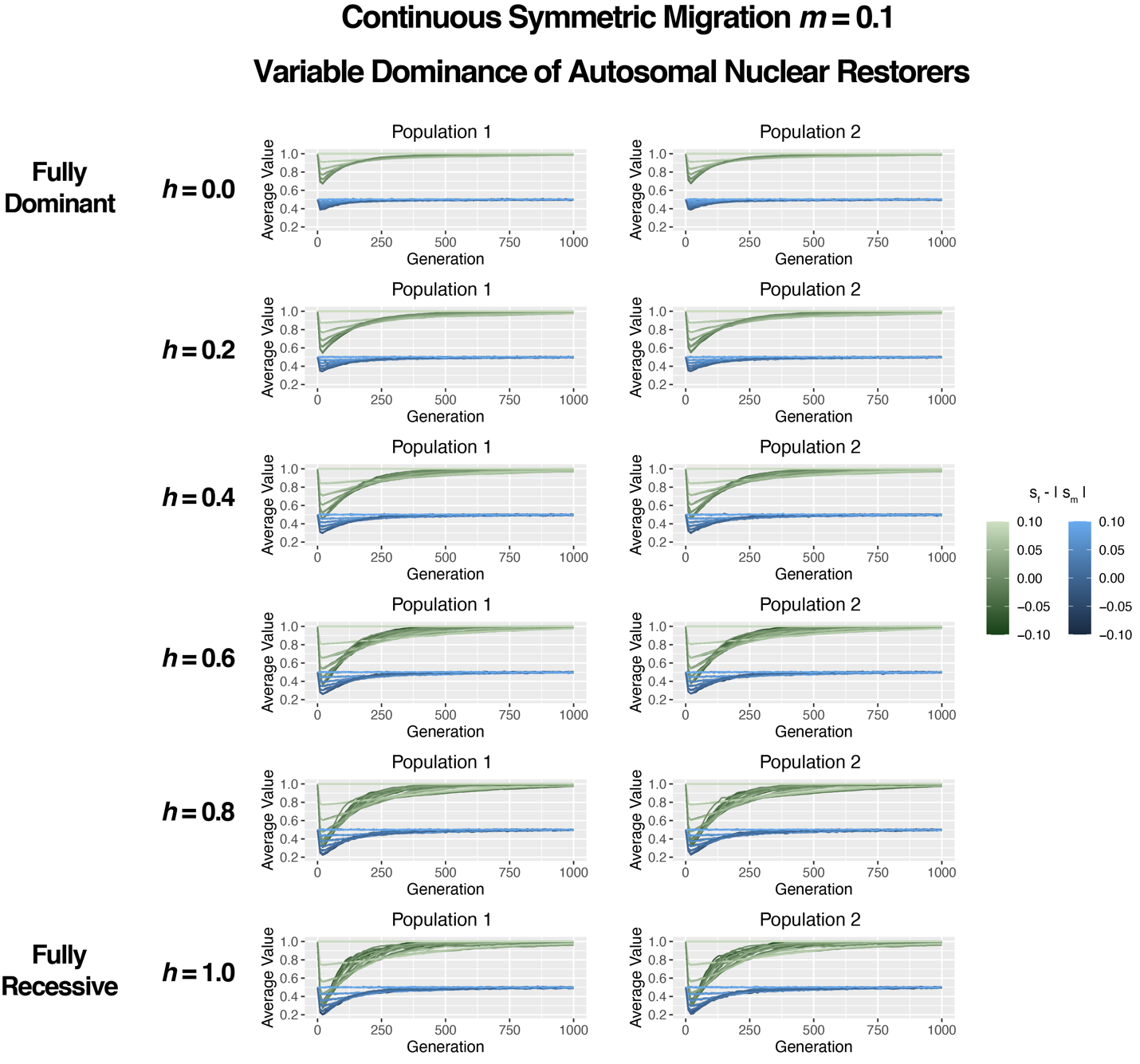

**Figure S10. Mean Male Fitness and Sex Ratio Trajectories for population 1 (left) and population 2 (right) under continuous symmetric migration rates *m* = 0.1 for Autosomal Restorers if we vary the dominance effect of autosomal nuclear restorers (each row).** Mean male fitness trajectories are colored in green, while the sex ratio is colored in blue. The specific color refers to the magnitude of difference between the advantage of Mother’s Curse variants in females and the deleterious effect of these variants in males. Given how we set up fitness evaluations, *h* = 0.0 represents a fully dominant restorer, while *h* = 1.0 represents a full recessive restorer.

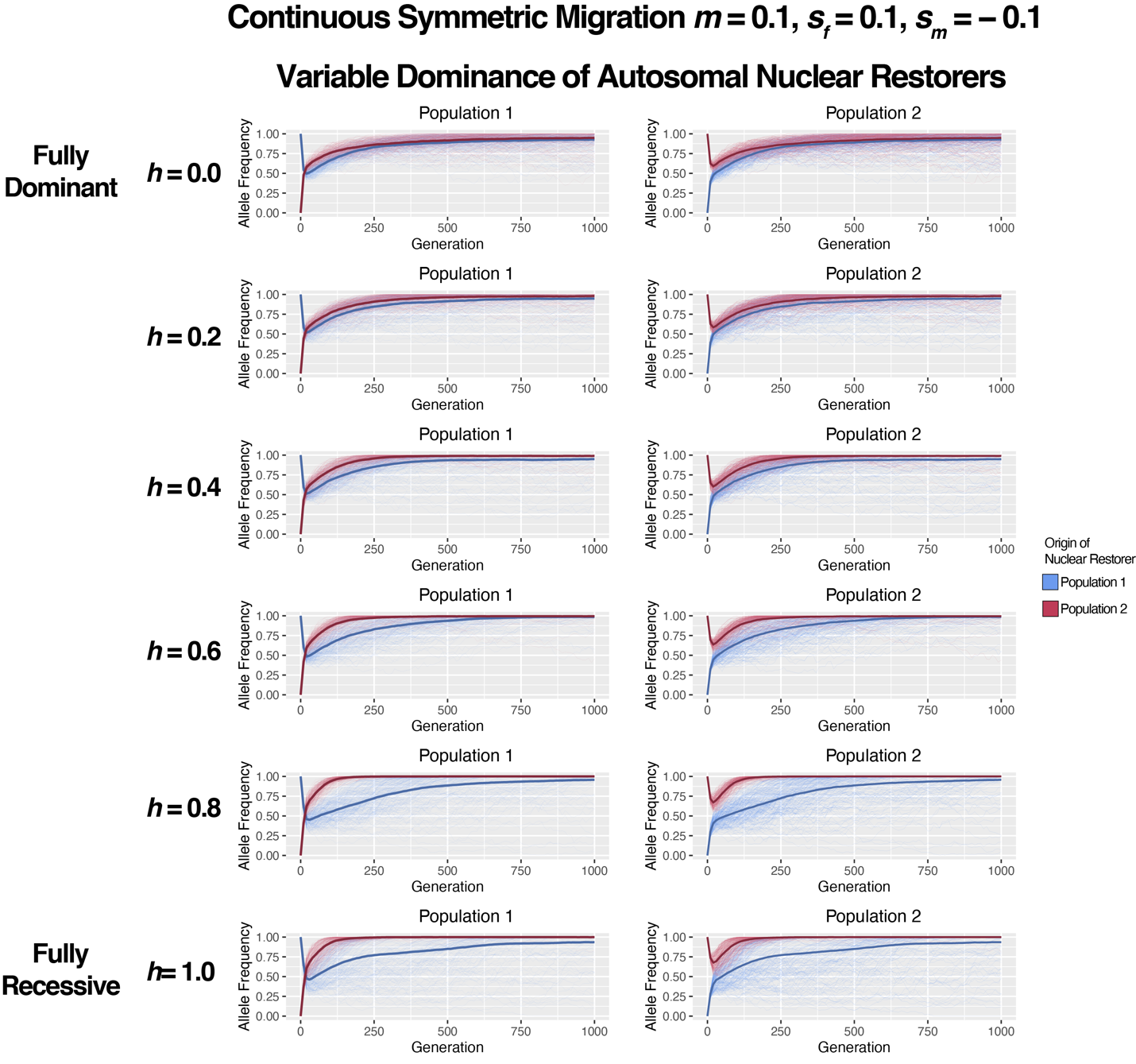

**Figure S11. Allele Frequency Trajectories for both sets of nuclear restorers, those that originated in population 1 (blue) and population 2 (magenta) in each population (left and right) continuous symmetric migration rate *m* = 0.1, *s_f_* = 0.1, and *s_m_* = −0.1 for Autosomal Restorers if we vary the dominance effect of autosomal nuclear restorers (each row)**. Each line represents the trajectory of a specific restorer. The darker, bold line represents the mean across all trajectories for that class of nuclear restorers. The first row shows autosomal restorers, the second row shows X-linked restorers, and the third row shows Y-linked restorers. Given how we set up fitness evaluations, *h* = 0.0 represents a fully dominant restorer, while *h* = 1.0 represents a full recessive restorer.

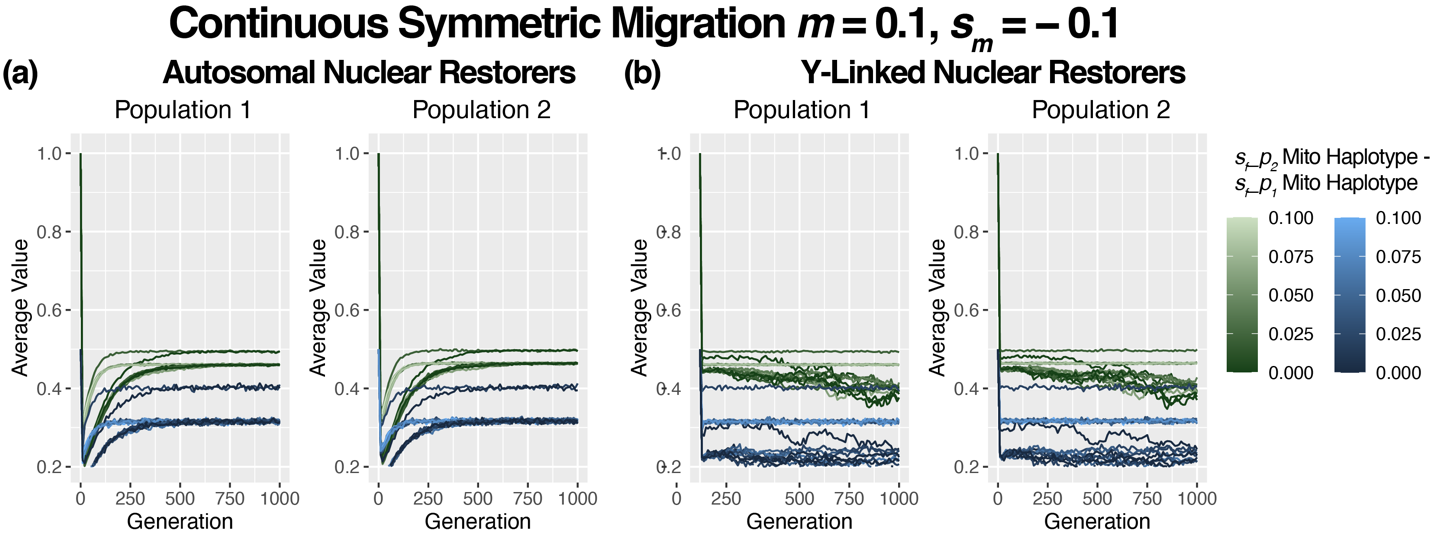
**Figure S12. Mean Male Fitness and Sex Ratio Trajectories for population 1 (left) and population 2 (right) under Continuous Symmetric Migration rate *m* = 0.1 for Autosomal (a) and Y-linked Nuclear Restorers (b) if we assign different fitness in females for each mitochondrial haplogroup.** The mitochondrial haplogroup native to population 1 has *s_f_* = 1.0 in females, while the haplogroup native to population 2 has *s_f_* = 1.1.Mean male fitness trajectories are colored in green, while the sex ratio is colored in blue. The specific color refers to the magnitude of difference between the two advantage of Mother’s Curse variants for a fixed deleterious effect in males (*s_m_* = −0.1**)**.

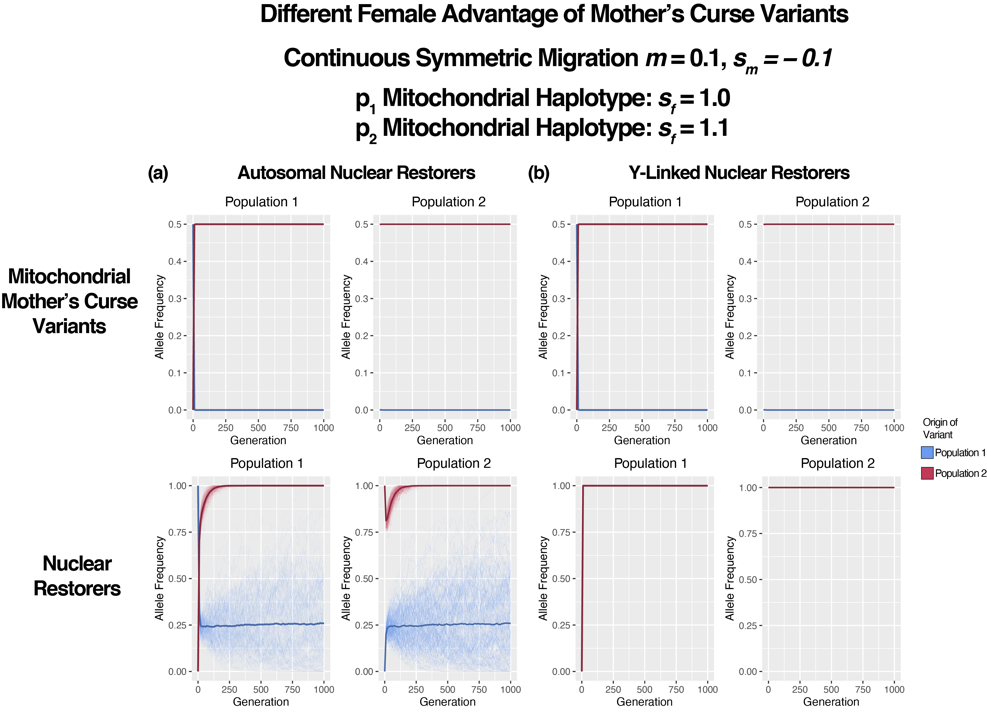

**Figure S13. Allele Frequency Trajectories for both sets of Mother’s Curse variants (Top Row) and nuclear restorers (Bottom Row), those that originated in population 1 (blue) and population 2 (magenta) in each population (left and right) under continuous symmetric migration rate *m* = 0.1 for Autosomal (a) and Y-linked Nuclear Restorers (b) if we assign different fitness in females for each mitochondrial haplogroup.** The mitochondrial haplogroup native to population 1 has *s_f_* = 1.0 in females, while the haplogroup native to population 2 has *s_f_* = 1.1. Each line represents the trajectory of a specific restorer. The darker, bold line represents the mean across all trajectories for that class of nuclear restorers. This confirms that using differential female fitness results in population replacement

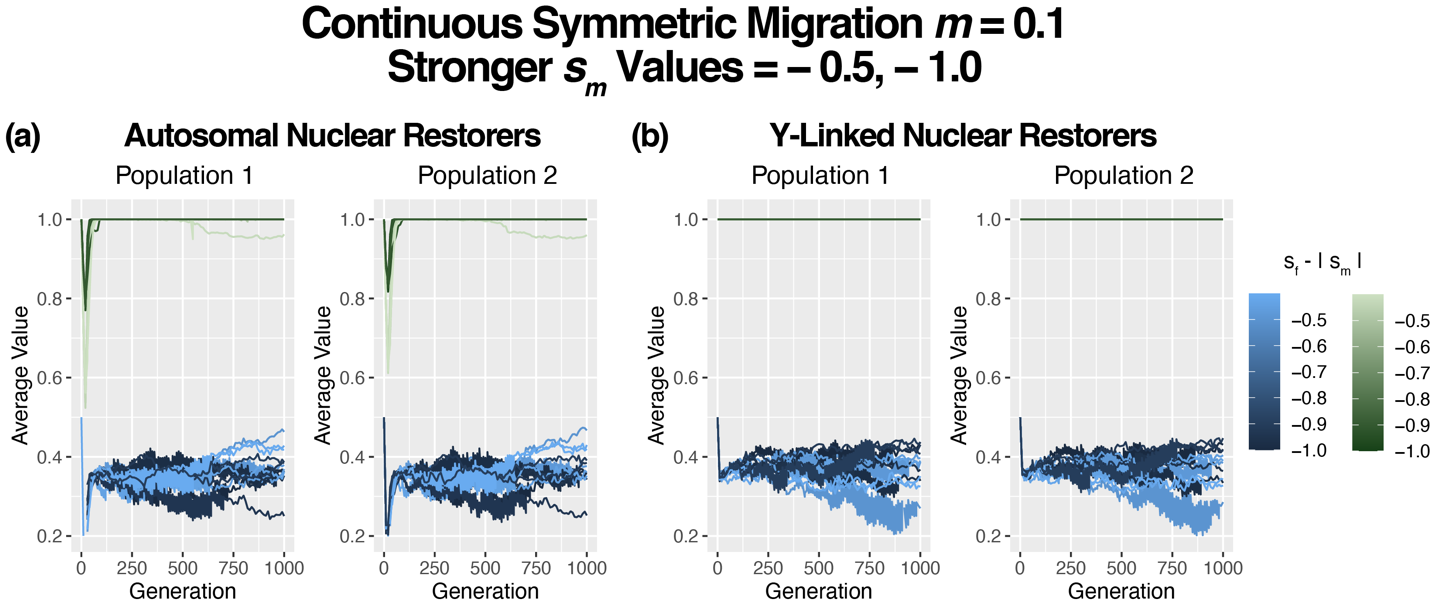

**Figure S14. Mean Male Fitness and Sex Ratio Trajectories for population 1 (left) and population 2 (right) under Continuous Symmetric Migration rate *m* = 0.1 for Autosomal (a) and Y-linked Nuclear Restorers (b) if we use more extreme values for *s_m_*.** Mean male fitness trajectories are colored in green, while the sex ratio is colored in blue. The specific color refers to the magnitude of difference between the advantage of Mother’s Curse variants in females and the deleterious effect of these variants in males.

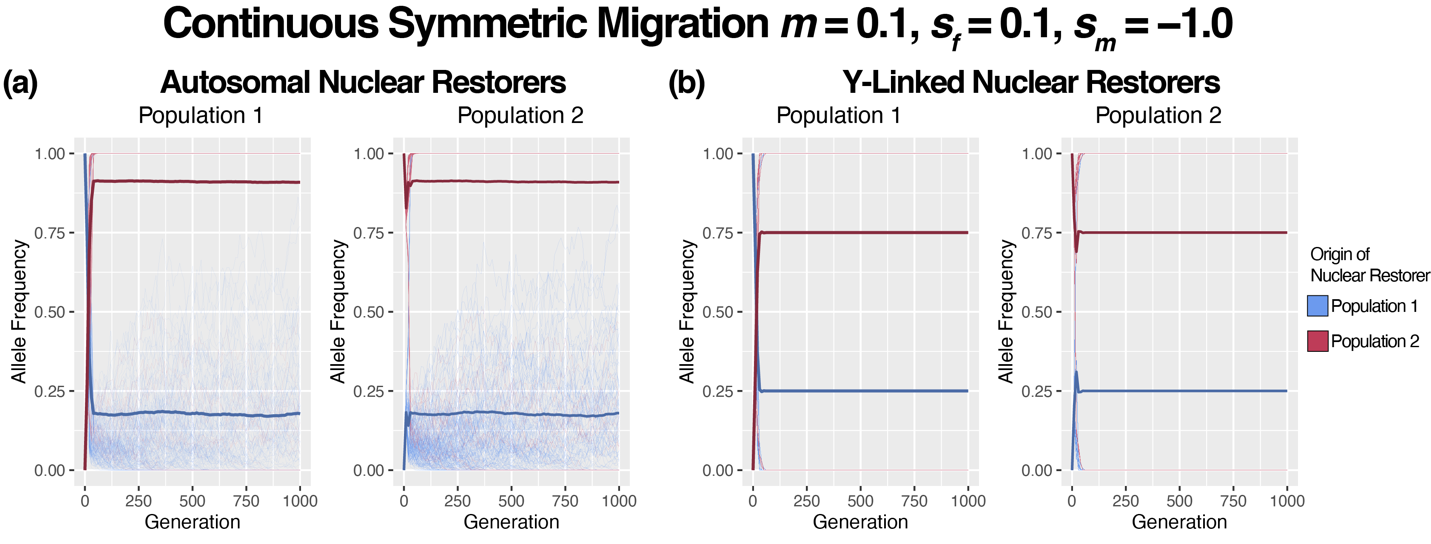
**Figure S15. Allele Frequency Trajectories for both sets of nuclear restorers, those that originated in population 1 (blue) and population 2 (magenta) in each population (left and right) under continuous symmetric migration rate *m* = 0.1, *s_f_* = 0.1, and *s_m_* = -1.0 for Autosomal (a) and Y-linked Nuclear Restorers (b) if we use more extreme values for *s_m_*.** Each line represents the trajectory of a specific restorer. The darker, bold line represents the mean across all trajectories for that class of nuclear restorers.
